# Fecal virome transfer improves proliferation of commensal gut *Akkermansia muciniphila* and unexpectedly enhances the fertility rate in laboratory mice

**DOI:** 10.1101/2022.06.30.498243

**Authors:** Torben Sølbeck Rasmussen, Caroline M. Junker Mentzel, Malene Refslund Danielsen, Rasmus Riemer Jakobsen, Line Sidsel Fisker Zachariassen, Josue Leonardo Castro Mejia, Lars Hestbjerg Hansen, Axel Kornerup Hansen, Dennis Sandris Nielsen

## Abstract

Probiotics have been suggested as nutritional supplements to improve gastrointestinal health. However, the probiotics marketed today only colonize the densely populated gut to a limited extent. Bacteriophages comprise the majority of viruses in the human gut virome and there are strong indications that they play important roles in shaping the gut microbiome (GM). Here we investigate the use of fecal virome transplantation (FVT) as a mean to alter GM composition to lead the way for persistent colonization of two types of probiotics: *Lacticaseibacillus rhamnosus* GG (LGG) representing a well-established probiotic and *Akkermansia muciniphila* (AKM) representing a putative next-generation probiotic. Male and female C57BL/6NTac mice were cohoused in pairs at 4 weeks of age and received the following treatment by oral gavage at week 5 and 6: AKM+FVT, probiotic sham (Pro-sham)+FVT, LGG+Saline, AKM+Saline, and control (Pro-sham+Saline). The FVT originated from donor mice with high relative abundance of *A. muciniphila*. All animals were terminated at age 9 weeks. The FVT treatment did not increase the relative abundance of the administered LGG or AKM in the recipient mice. Instead FVT significantly (p<0.05) increased the abundance of naturally occurring *A. muciniphila* compared to the control. This highlights the potential of stimulating the commensal “probiotics” that already are permanent members of the gut. Being co-housed male and female, a fraction of the female mice became pregnant. Unexpectedly, the FVT treated mice were found to have a significantly (p<0.05) higher fertility rate independent of probiotic administration. These preliminary observations urge for follow-up studies investigating GM/fertility interactions.

## Introduction

During the last decade it has become commonly accepted that gut microbiome (GM) imbalances (dysbiosis) play important roles in the etiology of a number of diseases [1–3]. Probiotics has been suggested as a tool to restore GM balance [4,5] and are defined as live microorganisms that when ingested in adequate amounts confer a health benefit to the host [6]. However, traditional probiotics, mainly lactobacilli and bifidobacteria, in general have no or only modest influence on GM composition [7]. So-called next generation probiotics, like *Akkermansia muciniphila*, have recently been suggested for alleviating GM-associated malfunctions [8,9]. Persistent beneficial effects are challenged by the difficulties of the administered bacteria to become a permanent and adequately abundant member of the densely populated GM [10,11].

Mounting evidence suggests that the gut viral community plays a pivotal role in shaping the composition of the GM [12,13]. The gut virome is predominated by prokaryotic viruses [14], including bacteriophages (phages), which are viruses that attack bacteria in a host-specific manner [15]. A transfer of sterile filtered feces (containing phages, but no intact bacterial cells) from a healthy donor have shown to successfully treat recurrent *Clostridioides difficile* infections (rCDI) in human recipients [16]. Other studies using sterile filtered feces have reported to alleviate symptoms of type-2-diabetes (T2D) and obesity in mice [17], and to prevent the development of necrotizing enterocolitis [18] in preterm piglets. These changes in phenotype may be driven by a phage-mediated modulation of the GM [17–21]. In all cases, when transferring the fecal viral components, a significant change in the bacterial diversity and composition was observed, with the bacterial GM-component of the recipients becoming more like the GM of the donors. We will refer to this approach as fecal virome transplantation (FVT).

Using co-housed male and female laboratory mice as model, we hypothesized that initial phage-mediated disturbance of the existing bacterial landscape in the GM (using FVT) would improve the enteric engraftment and abundance of the administered probiotic bacteria (*Lactocaseibacillus rhamnosus* GG or *A. muciniphila*). We did the following considerations to maximize our chances for evaluating a successful enteric engraftment of *A. muciniphila*: (i) C57BL/6NTac (B6N mice from Taconic) mice were selected since previous experience have shown a low relative abundance of *A. muciniphila* in mice from this vendor [22], (ii) the *A. muciniphila* YL-44 strain was used in this study due to its enteric origin from the genetically close related C57BL/6J (B6J) wildtype mouse, and (iii) the FVT virome represented a gut virome from mice donors with relatively high *A. muciniphila* abundance [22].

## Results

Here we investigated the potential of transferring an *A. muciniphila* rich GM phenotype via fecal virome transplantation (FVT) from lean mouse donors to lean recipients to improve persistent colonization of two probiotics; *L. rhamnosus GG* (LGG) or *A. muciniphila* (AKM). Fecal samples from three different timepoints were included to investigate the level of probiotic engraftment and GM changes over time: baseline, 6 days after 2^nd^ intervention, and at termination. See Figure 1 for the experimental design of the animal model.

**Figure 1:**
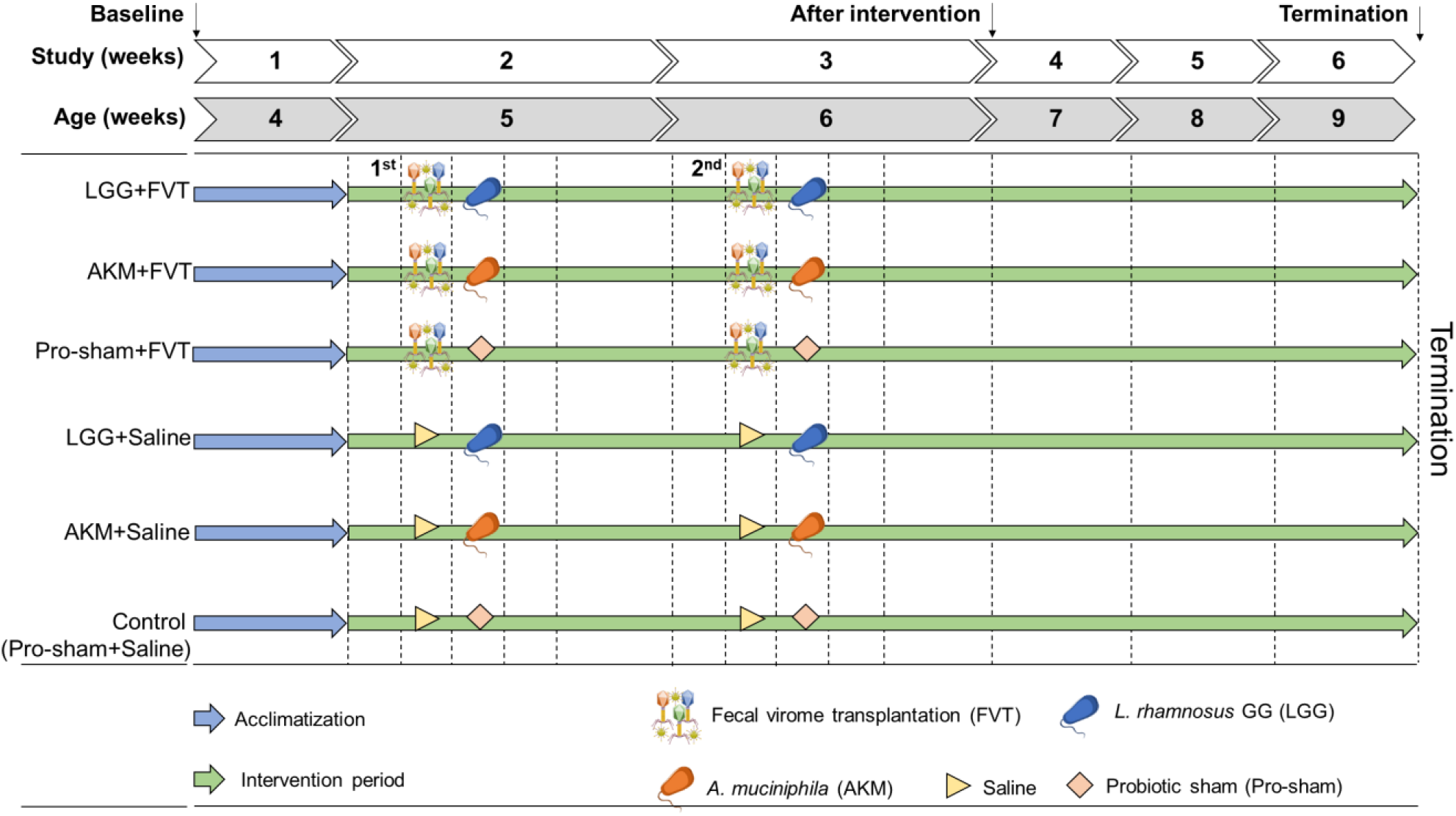
Experimental setup of the animal model. 24 male and 24 female C57BL/6NTac mice (4 weeks old) were divided into six groups: LGG+FVT, AKM+FVT, Pro-sham+FVT, LGG+Saline, and AKM+Saline, control (Pro-sham+Saline). The Saline consisted of SM buffer and Pro-Sham of Intralipid®. The mice were administered 1 M sodium bicarbonate prior oral gavage of FVT/Saline to protect the viral community against the acidic environment in the stomach. The day after, the mice were inoculated with probiotic solutions of LGG/AKM/Pro-sham suspended in Intralipid® which constituted the 1^st^ inoculation. The same procedure was repeated as the 2^nd^ inoculation one week after. The mice were fed *ad libitum* low-fat diet (LFD) for the entire study (6 weeks) until termination at age 9 weeks. Fecal samples from baseline, after intervention, and termination were analyzed in this study. Abbreviations: *Lacticaseibacillus rhamnosus* GG = LGG, *Akkermansia muciniphila* = AKM, fecal virome transplantation = FVT, Pro-sham = probiotic sham.

### FVT enhanced the abundance of natural occurring Akkermansia muciniphila strains

Our hypothesis was that initial disruption of the GM landscape driven by the FVT would lead to an increase in the abundance of AKM and/or LGG after probiotic administration. However, we did not observe any significant effect of the FVT on the AKM/LGG abundance after intervention nor at termination (Figure 2). Instead, we observed at termination that FVT had increased (p < 0.05) the abundance of, what would be expected to be, naturally occurring (native) *A. muciniphila* strains in mice that were not provided AKM as probiotic compared to mice neither provided FVT nor AKM (Figure 2A & Figure 2C). The abundance of *A. muciniphila* strains at baseline were significantly lower (p < 0.05) compared to termination in AKM+FVT and Pro-sham+FVT mice, while tending lower (p < 0.1) in the LGG+FVT mice as well (Figure S1).

**Figure 2:**
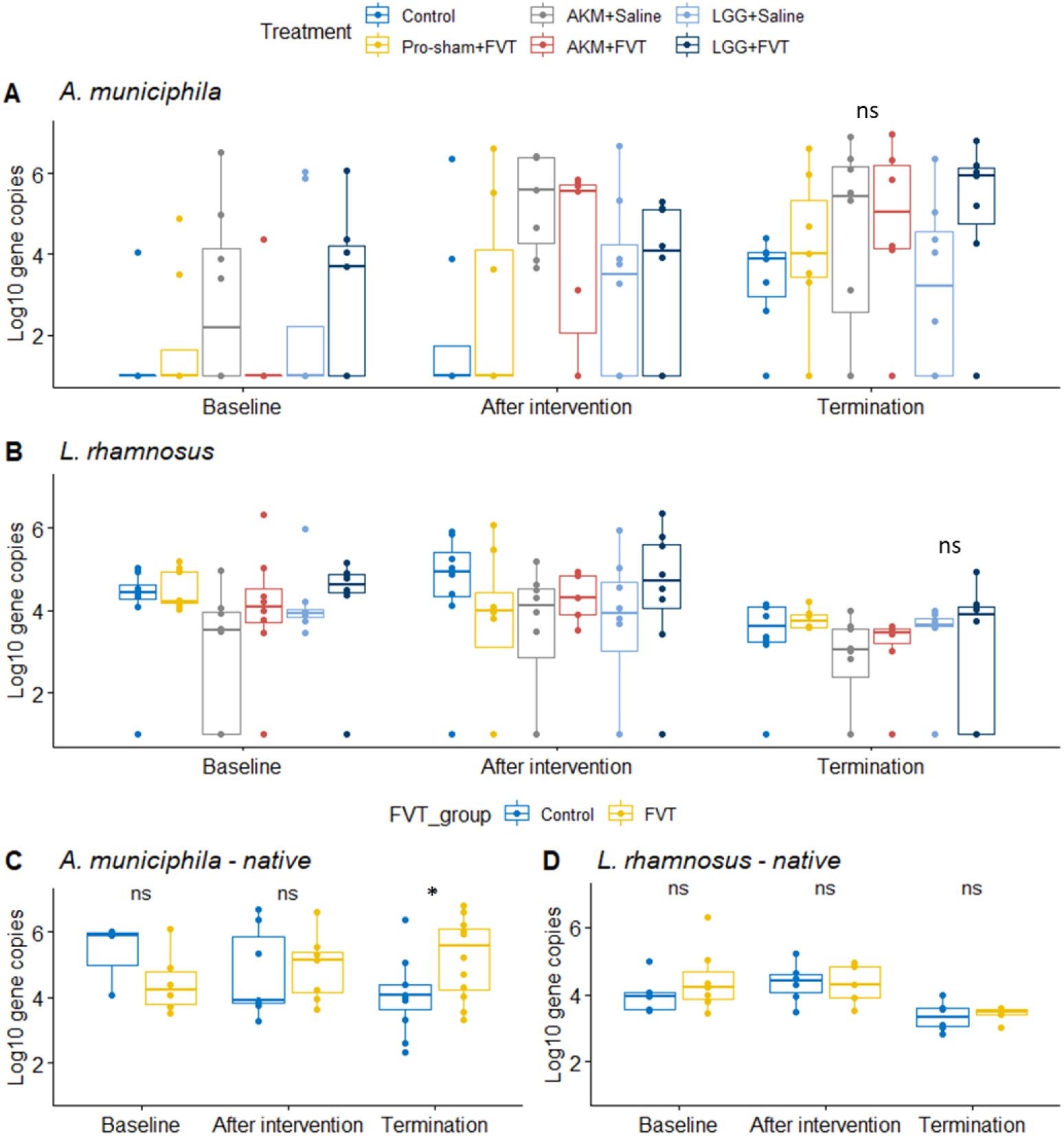
qPCR with *L. rhamnosus* and *A. muciniphila* specific primers were used to assess the abundance of gene copies per gram feces over the time span of baseline, after intervention, and termination. A) The development of the *A. muciniphila* abundance, B) *L. rhamnosus* abundance. C) The abundance development of native *A. muciniphila* strains for mice receiving FVT and no AKM (Pro-sham+FVT and LGG+FVT) compared to not receiving FVT or AKM (LGG+Saline and control). D) The abundance development of native *L. rhamnosus* strains for mice receiving FVT and no LGG (Pro-sham+FVT and AKM+FVT) compared to not receiving FVT or LGG (AKM+Saline and control). Abbreviations: *Lacticaseibacillus rhamnosus* GG = LGG, *Akkermansia muciniphila* = AKM, fecal virome transplantation = FVT, Pro-sham = probiotic sham.

This indicated that the FVT had improved the growth conditions of naturally occurring *A. muciniphila* strains due to yet unknown environmental changes. The AKM+Saline mice had similar *A. muciniphila* abundance as the FVT groups. Additional experiments were performed to rule out that the FVT initially contained any *A. muciniphila* strains (Figure S2). The sterile filtered donor feces (used for FVT) were incubated in 96 hours on GAM agar plates from which eight colonies appeared. Cell morphology of these colonies was imaged with phase-contrast microscopy, and subsequently screened with *A. muciniphila* specific primers in both a PCR and qPCR assay. Neither colony morphology, cell morphology, PCR nor qPCR indicated any traces of *A. muciniphila* in the applied donor FVT (Figure S2).

### FVT leads to a reduction in bacterial diversity of the GM component

FVT significantly (p < 0.05) decreased the bacterial Shannon diversity index in the LGG+FVT and Pro-sham+FVT mice at termination (9 weeks of age) when compared with the AKM+Saline, LGG+Saline, and the control mice (Figure 3A). Whereas the Shannon diversity index of the AKM+FVT mice remained unchanged compared to the control mice, hence suggesting that AKM may have counteracted the decrease in the Shannon diversity that was associated to the FVT treatment (Figure 3A). The initial bacterial diversity at baseline was similar between all groups (Figure 2A). The most abundant genus in all groups at all time-points were *Lactobacillus* (Figure S3). The FVT-associated differences in the bacterial Shannon diversity index were not reflected in the bacterial composition analysis (Bray-Curtis dissimilarity), since no significant differences were observed between treatments at all three timepoints (Figure 3B). Probably due to the state of pregnancy, the sex of the animals (male vs female) showed significant (p < 0.001) differences in their bacterial composition at termination (Figure S4). Differential abundance (DA) analysis showed that *Candidatus* Arhtromitus (segmented filamentous bacteria) and *A. muciniphila* amplicon sequence variants (ASVs) were significantly (p < 0.05) increased in relative abundance at termination in mice receiving FVT compared to the other groups (Figure 3C). Thus, clearly supporting the FVT-mediated enhancement of *A. muciniphila* abundance measured by the qPCR analysis. The administration of AKM significantly increased (p < 0.05) the relative abundance of *Ruminococcus gnavus* (Figure S5) but not *per se* influence *A. muciniphila* relative abundance in the recipients.

**Figure 3:**
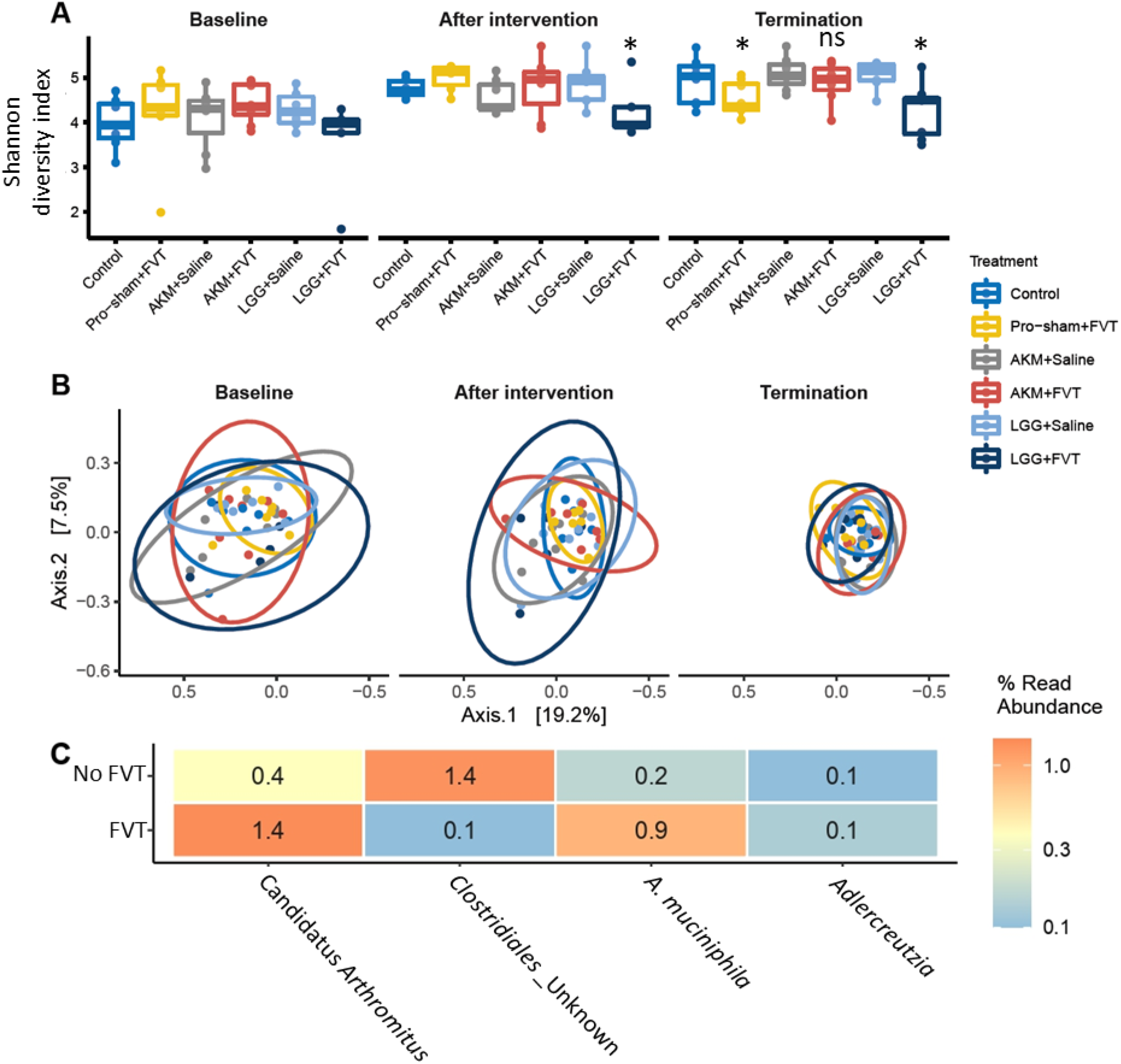
Gut bacteriome analysis of A) the bacterial diversity (Shannon diversity index) and B) PCoA plots of the bacterial composition (Bray Curtis dissimilarity) at baseline, after intervention, and termination. C) Differential abundance analysis of ASVs with significant (p < 0.05) different relative abundance between FVT treated mice and mice not receiving FVT. Abbreviations: *Lacticaseibacillus rhamnosus* GG = LGG, *Akkermansia muciniphila* = AKM, fecal virome transplantation = FVT, Pro-sham = probiotic sham, ASV = amplicon sequence variant.

### Probiotic and FVT intervention may have changed the viral GM profile

The viral Shannon diversity index at termination (9 weeks of age) was affected by FVT (p = 0.032) as well as the administration of the probiotics AKM (tendency, p = 0.07) and LGG (p = 0.006) when compared to the control mice (Figure 4A and Figure S6). The effects of AKM (p = 0.025) and LGG (p = 0.014) were also reflected on the viral composition at termination (Figure S7). The sex of the animals appeared to influence (p < 0.05) the viral community composition across all time points (Figure S4). The donor FVT virome consisted of more than 90% *Microviridae* viruses and was markedly different in both viral diversity and composition (Figure 4A & Figure 4B) compared to the recipient gut virome (Figure S8). DA analysis was performed using both the predicted bacterial hosts and raw viral taxonomy at termination (9 weeks of age). These analyzes showed a significant increase in the relative abundance of predicted hosts belonging of the taxa *Lachnospiraceae, Parabacteroides*, and *Bacteroides* in FVT treated mice (Figure S9), and an increase in the relative abundance of *Petitvirales* (likely *Microviridae*) when comparing with mice not receiving FVT (Figure 4C). It could be speculated that the elevated level of *Petitvirales* in the FVT treated mice was driven by the highly *Microviridae* abundant (> 90% of relative abundance) donor virome.

**Figure 4:**
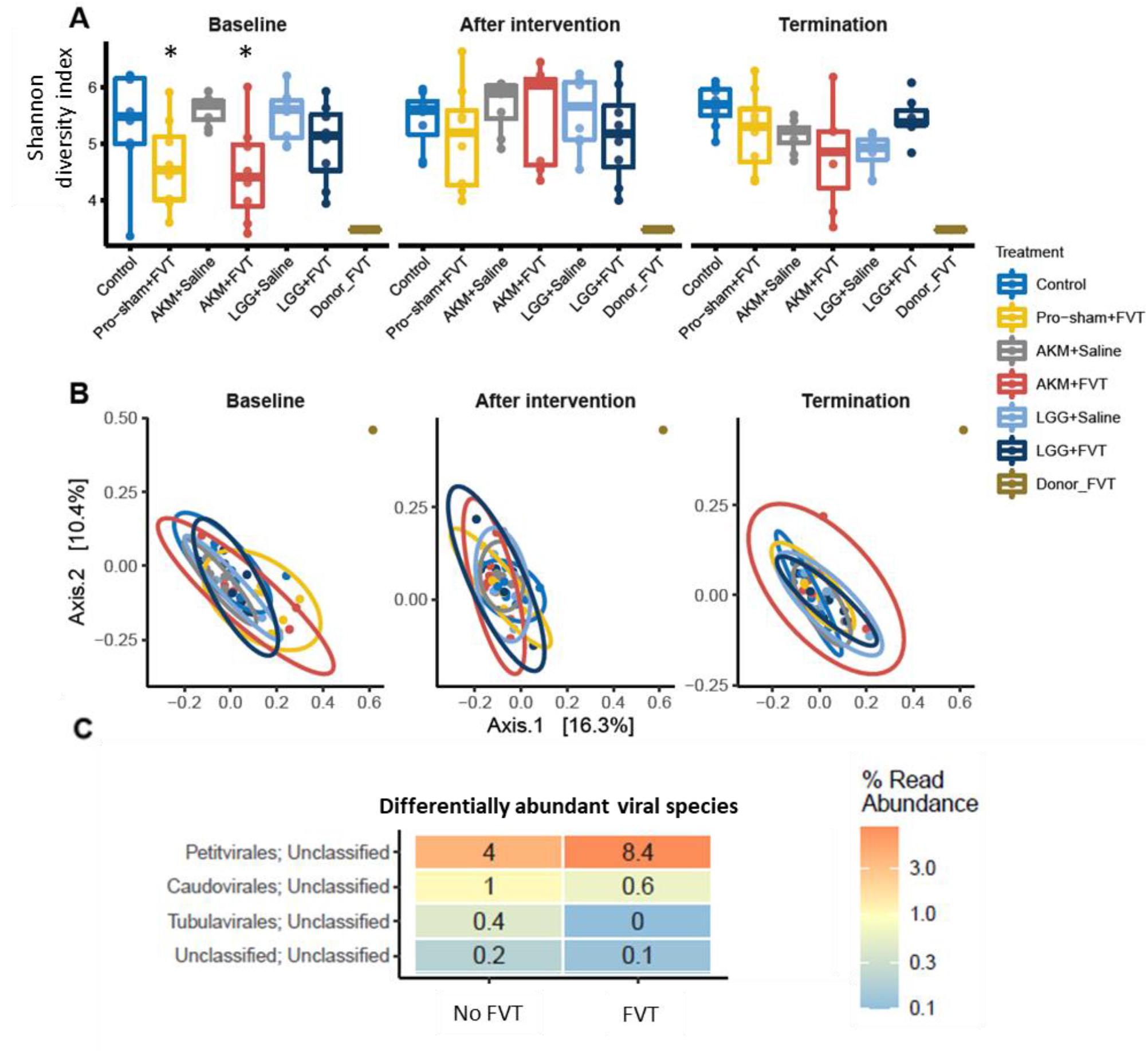
Gut virome analysis of A) the viral diversity (Shannon diversity index) and B) PCoA plots of the viral composition (Bray Curtis dissimilarity) at baseline, after intervention, and termination. C) Differential abundance analysis of viral contigs with significant (p < 0.05) different relative abundance between FVT treated mice and mice not receiving FVT. Abbreviations: *Lacticaseibacillus rhamnosus* GG = LGG, *Akkermansia muciniphila* = AKM, fecal virome transplantation = FVT, Pro-sham = probiotic sham.

The viral Shannon diversity at baseline of the AKM+FVT and Pro-sham+FVT mice were significantly lower (p < 0.05) compared to control (Figure 4A), while the diversity of the remaining treatment groups was similar to the control mice. This initial variance also tended to be reflected (p < 0.084) on the viral composition at baseline, but these differences were diminished at termination (Figure 4B). The extent to which the above-mentioned inter-group differences were associated with the baseline variance of viral diversity and composition was not clear.

### A. muciniphila affects expression of a gene involved in mucin-production and limits an inflammatory response associated to FVT

The ileum tissue was investigated for changes in the expression levels of genes associated to inflammatory responses. Interestingly, mice with the highest abundance at termination of the mucin-degrading *A. muciniphila* (Figure 2A) had either significantly (p < 0.05, AKM+Saline and AKM+FVT) or tended towards (p < 0.1, LGG+FVT) lowered gene expression of genes related to inflammatory responses compared to control mice (Figure 5A). The *Muc1* gene is involved in mucin production [23]. Whether this affected the mouse phenotype was not clear. The expression of nine genes that are involved in inflammation and as a response on infection (*Clc2, Ccr10, Ctla4, Cxcl1, Il1b, Il4*, Il6, *Retnlb, Timp1)*, were significantly (p < 0.05) elevated in Pro-sham+FVT compared to control mice (Figure 5A - 5J). Additionally, two genes (*Ffar2* and *Ffar3*) involved in both energy homeostasis and intestinal immunity were respectively increased (tendency, p = 0.06) or decreased (p = 0.01) in the Pro-sham+FVT compared to control mice (Figure 5K & 5L). Excluding the Pro-sham+FVT mice, the *nod2* gene was lowered in expression in all treatment groups compared to control (Figure 5M).

**Figure 5:**
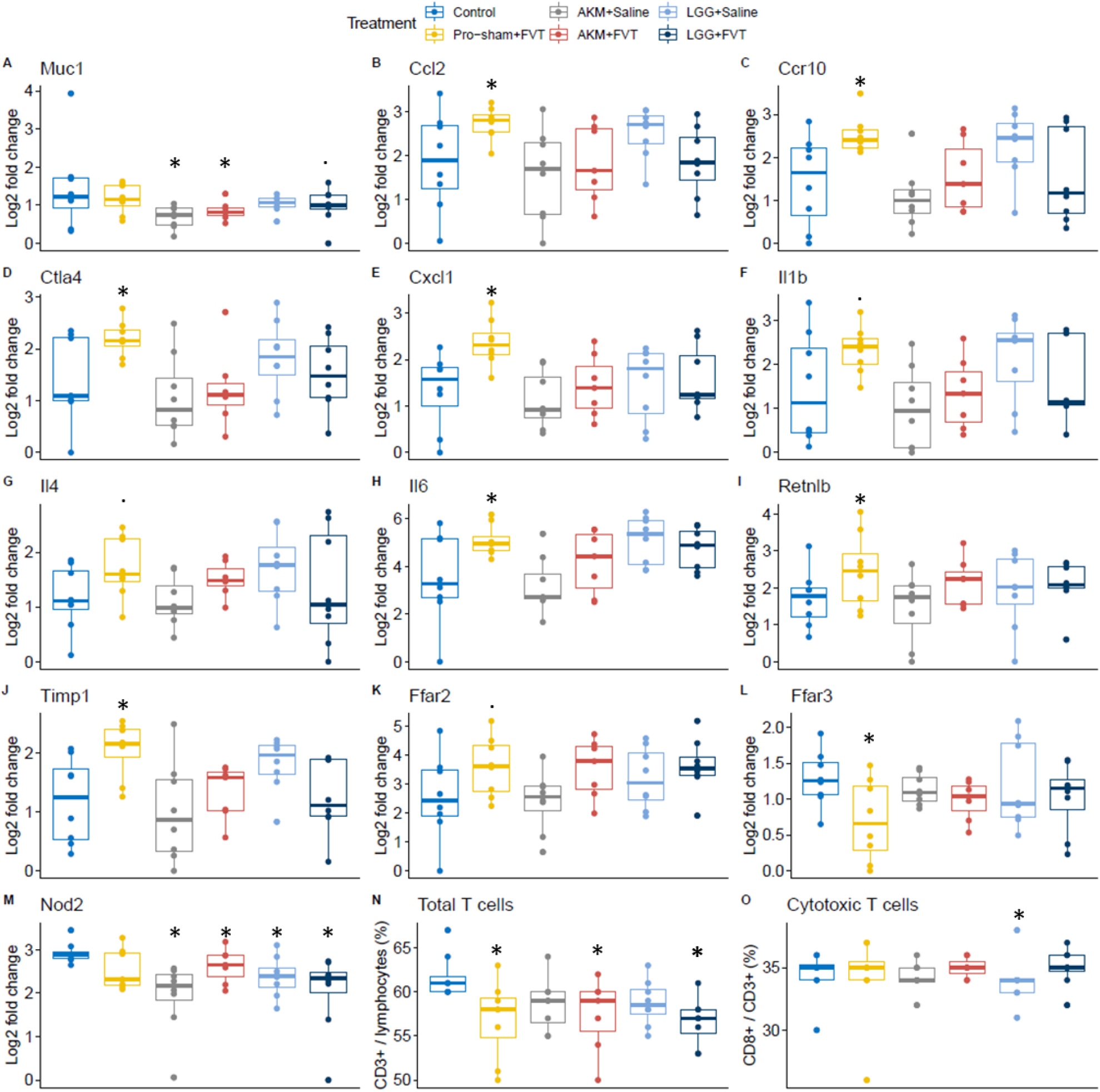
The ileum tissue was investigated for changes in the expressions levels of genes associated to inflammatory responses at termination. In A) to M) the expression levels of these selected genes are shown, and the different treatment groups are compared with the control mice. The relative abundance of total number of T cells N) and cytotoxic T cells (CD8+ T cells) P) were measured. Abbreviations: “*” = (p < 0.05), “.” = (p < 0.1), *Lacticaseibacillus rhamnosus* GG = LGG, *Akkermansia muciniphila* = AKM, fecal virome transplantation = FVT, Pro-sham = probiotic sham.

Altogether, these changes in gene expression indicate that the FVT of the Pro-sham+FVT mice had initiated an inflammatory response, possibly mediated by the presence of eukaryotic and/or prokaryotic viruses that were transferred with the FVT. The administration of AKM along with FVT seemed to counteract this inflammatory response, since the expression of none of the above-mentioned 11 genes were changed in the AKM+Saline or AKM+FVT compared to control mice (Figure 5A – 5L).

Fluorescence-activated cell sorting (FACS) was performed to evaluate the presence of selected immune cells in the mesenteric lymph node (MLN) at termination. The FVT treated male and female mice expressed a significant (p = 0.043) decrease in the total number of T cells (CD3^+^ lymphocytes) (Figure 5N), while mice provided only LGG had a significant decrease (p = 0.01) in the level of cytotoxic T cells (CD8^+^/CD3^+^) (Figure 5O). The level of cytotoxic T cells in the MLN was not affected by the FVT treatment, suggesting that the increased expression of inflammatory genes in the ileum tissue, isolated from the Pro-sham+FVT mice, was not a systemic response. The levels of the remaining measured immune cells (CD11c, CD86^+^CD11c, CD11b^+^CD11c, CD103^+^CD11c) were not significantly affected by the sex of the animals, probiotics (AKM/LGG) or FVT.

### Increased fertility rate following FVT

The pregnancy status and fertility rate (number of fetuses or born pups) were evaluated for each female mouse (Figure 6 and Figure S10) due to the natural consequences of pairing male and female mice in cages. Surprisingly, the FVT treated female mice (Pro-sham+FVT, AKM+FVT, and LGG+FVT) exhibited a significant increase in both fertility rate (p = 0.014, Figure 6A) and pregnancy status (p = 0.025), Figure 6B) compared to controls. These observations were independent of the administered probiotics LGG/AKM. It was also observed that none of the female AKM+Saline mice (n = 4) were pregnant (Figure S10).

**Figure 6:**
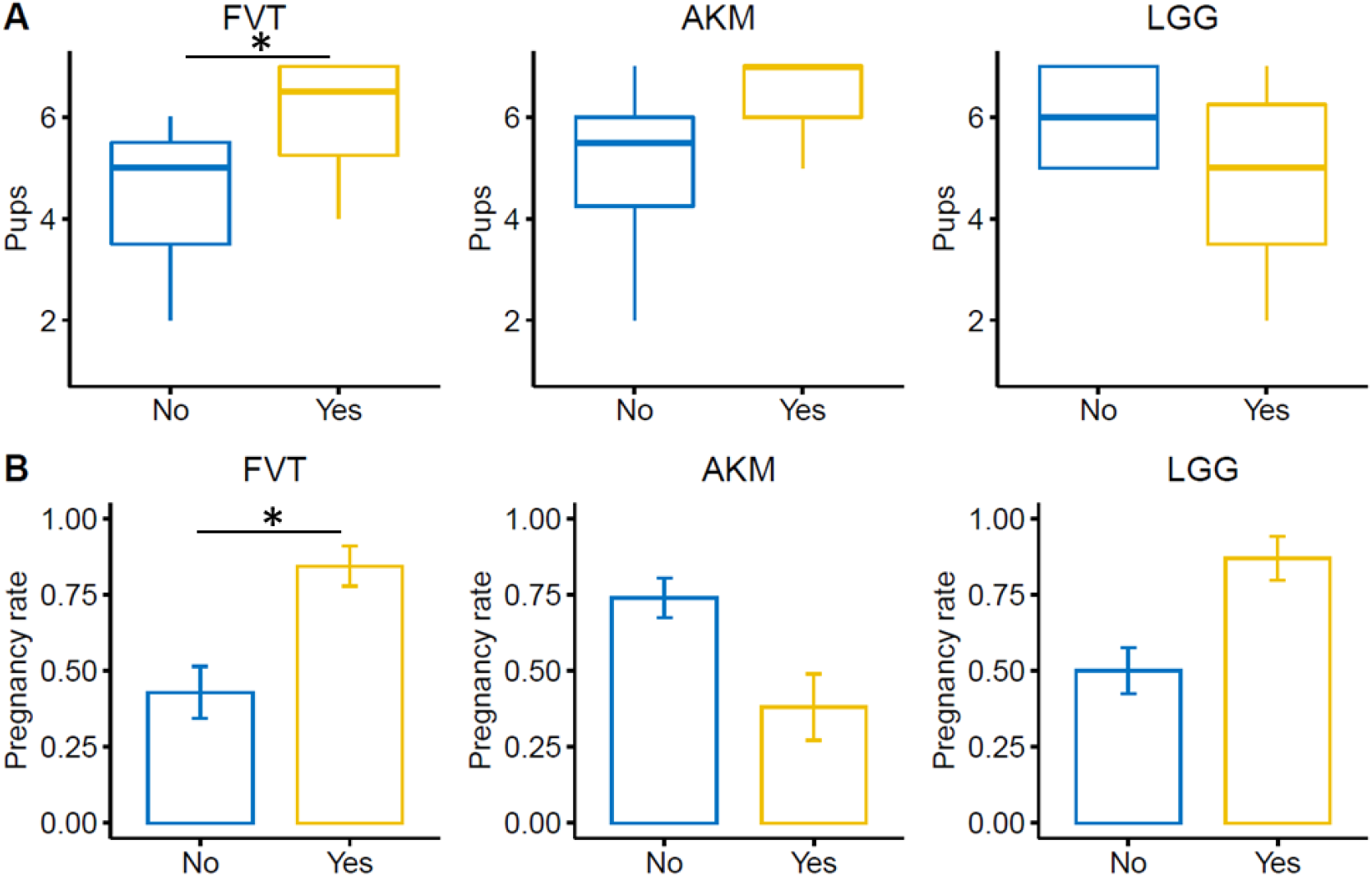
Bar plots of the fertility rate (number of pups) and pregnancy rate. A) The observed number of pups (born or as fetuses) based on a linear model (y∼FVT+probiotics). B) The mean distribution of the of the binary event of pregnancy. Only the 23 female mice that received either FVT/Saline along with probiotic solutions of AKM/LGG/Pro-sham was included in the statistical analysis that was based on a generalized logistic regression model (Figure S10). Abbreviations: *Lacticaseibacillus rhamnosus* GG = LGG, *Akkermansia muciniphila* = AKM, fecal virome transplantation = FVT, Pro-sham = probiotic sham, “*” = (p < 0.05).

## Discussion

Here we investigated the potential of using phage-mediated “GM disturbance” to improve the engraftment of two different probiotics (*A. muciniphila;* AKM and *L. rhamnosus GG;* LGG) in lean mouse recipients over a time span of 5 weeks. The hypothesis was that fecal virome transplantation (FVT) followed by probiotic administration would increase the chance of persistent enteric colonization of the probiotic bacteria. However, the results did not support the hypothesis. Instead, specie specific qPCR analysis showed that the FVT treatment at termination (9 weeks of age) had significantly (p < 0.05) increased the abundance of naturally occurring *A. muciniphila* strains, compared to non-FVT treated mice (Figure 2A & 2C). We ruled out that other *A. muciniphila* strains were present in the applied FVT virome (Figure S2). The control (Pro-sham+Saline) and LGG+Saline mice also increased their *A. muciniphila* abundance over time which likely can be explained by regular GM maturation [24,25]. However, *A. muciniphila* abundance remained 1-2 log higher (p = 0.065) in the FVT/AKM groups at termination compared with the mice receiving neither AKM nor FVT (Figure S1). The applied FVT originated from a GM community that allowed a relative abundance of *A. muciniphila* > 6% [22]. Hence, the FVT-mediated enhancement of the abundance of naturally occurring *A. muciniphila* strains emphasize the potential of transferring a phenotype along with the FVT. A concept which has been reported in other studies as well [16–19]. The administered LGG did not persistently colonize regardless of FVT treatment (Figure 2).

The bacteriome analysis showed that FVT significantly (p < 0.05) decreased the bacterial Shannon diversity in the LGG+FVT and Pro-sham+FVT mice at termination compared to the control and the other treatment groups (Figure 3A). However, AKM may have counteracted this tendency since the bacterial diversity of the AKM+FVT was unaffected. The mechanism behind this observation is unknown. However, assuming that a phenotype can be transferred along with the FVT, the decrease in bacterial diversity may be associated to the very low viral diversity and markedly different viral composition in the transferred FVT donor virome (Figure 4A & 4B). It could therefore be hypothesized that a low viral diversity may also favor a low bacterial diversity, due to the inherit link between phages and their bacterial hosts. This may fit with a previous study where the viral diversity of the fecal donor virome was higher than the recipient virome and FVT resulted in increased bacterial diversity in the recipient mice compared to non-FVT treated mice [17]. More than 90% of the relative abundance of the FVT donor virome represented *Microviridae* viruses which was in accordance to the relative abundance of genetically identical B6N mice in a previous study [22]. Furthermore, PhiX (a *Microviridae* used to spike the metavirome sequencing) sequences were excluded from the analysis, thus the high relative abundance of *Microviridae* was not a technical artefact.

It should be noted that the initial viral diversity and composition at baseline appeared to vary within the treatment groups. The interpretation of changes in the viral diversity and composition were therefore challenged, although the analysis indicated that the FVT, AKM, and/or LGG may have affected (p < 0.05) both the viral Shannon diversity and composition irrespectively of initial group differences.

Both immune cell counting and the expression levels of genes involved inflammatory responses were measured to evaluate potential safety issues associated to the FVT from lean donors to lean recipients. Here especially the Pro-sham+FVT appeared with significantly (p < 0.05) increased gene expression levels compared to control mice (Figure 5) in the following pro-and/or anti-inflammatory related genes; *Ccl2* [26], *Ccr10* [27], *Ctla4* [28], *Cxcl1* [29], *Il1b* [30], *Il4* [31], *Il6* [32], *Retnlb* [33], *Timp1* [34], *Ffar2* [35,36], whereas *Ffar3* [35] and *Nod2* [37] decreased. Interestingly, the combination of AKM+FVT counteracted the elevated expression of these genes (Figure 5), which may be explained by previously suggested synergistic effects of combining probiotics and phages [38]. These indications of inflammatory response in the ileum tissue were not supported by immune cell counts in the mesenteric lymph node (MLN). The T cell counts were significantly lower (p < 0.05) in FVT treated mice compared to the control while the cytotoxic T cell and dendritic cell counts were similar when compared to control mice (Figure 5N & 5O). This suggest that the FVT did not activate a systemic immune response, but rather a more a local response due to the presence of foreign viral particles [39,40].

The administration of AKM in the AKM+Saline and AKM+FVT mice significantly decreased (p < 0.05) the expression of the *Muc1* gene that is a membrane-tethered mucin expressed on surfaces of epithelial cells – also in the intestine [23]. Both overexpression and knockout of the *Muc1* gene has been associated with the development of different cancer diseases [41–43], but have also been reported to be associated with anti-inflammatory effects by regulating toll-like receptor (TLR) expressions [44,45]. *A. muciniphila* is a mucin-degrading bacteria and have been suggested to have beneficial impact on human health, of which is linked to the regulation of the mucus thickness and gut barrier integrity [46]. Hence, it could be speculated that additional degradation of mucin by the administered AKM strain regulated the expression of *Muc1* to a neither over-expressive nor over-suppressive level.

The FVT treated mice were unexpectedly associated with a significantly (p < 0.05) increased fertility rate (Figure 6A) and pregnancy status (Figure 6B). The study was not designed to investigate fertility rates which also are reflected by the group sizes. Both the male and female mice were treated with FVT; thus, the basic premises of the experimental setup make it challenging to evaluate if the increased fertility rate and pregnancy status were due to improved sperm quality of the males and/or improved conditions for fertility in the female mice. Emerging evidence suggest that infertility should be added to the list of GM associated diseases [47–49], and the importance of validating this link to the GM is emphasized by infertility being estimated to affect up to 15% of couples world-wide [50,51]. Our observations are in line with other studies suggesting a link between fertility and the GM, e.g. the demonstrated improvement of spermatogenesis with FMT from healthy donors [52], as well as, impairment of spermatogenesis with FMT from donors with a dysbiotic GM [47]. In regard to females, links have been suggested between maternal obesity, gut dysbiosis, and inflammation [49]. New results have discovered a markedly increase in the abundance of *Bacteroides vulgatus* in the gut of polycystic ovary syndrome (PCOS) individuals, that through deconjugations of bile acids in the liver affects the interleukin-22 (IL-22) levels and ultimately the fertility [48]. Although additional experiments need to be conducted, it cannot be ruled out, that the transfer of AKM in the AKM+Saline mice led to similar cascading events that might have decreased the fertility of the male and/or female mice (Figure 6).

Conclusively, we here demonstrate that FVT increases the abundance of, what is expected to be, naturally occurring *A. muciniphila* strains in the recipient mice. The bulk and undefined nature of fecal viromes prevents any direct use as a commercial product. However, our results highlight the potential of using phage-mediated changes of the GM as a supplement to probiotics to enhance the growth of healthy commensals that outside the body are defined as probiotics. Furthermore, an unexpected event of increased fertility rate and pregnancy status was associated to the FVT treatment, which urge for additional studies specifically designed to clarify our observations.

## Methods

### Bacterial strains

The commercially available probiotic bacterium *Lacticaseibacillus rhamnosus* GG (LMG 18243, former *Lactobacillus rhamnosus* [53]) was included along with *Akkermansia muciniphila* YL-44 (DSM 26127), as a representative of next-generation probiotics [54].

### Preparation of inocula of L. rhamnosus (LGG) and A. muciniphila (AKM) for transfer to mice

*L. rhamnosus* GG (LGG) and *A. muciniphila* YL-44 (AKM) were both handled and incubated anaerobically as described previously [55]. AKM was incubated in Gifu Anaerobic Medium (GAM, HyServe, cat. no. 5422) and LGG in de Man Rogosa Sharpe broth (MRS, Merck, cat. no. 69966) in broth or agar plates containing 1.5% agar. In brief, GAM or MRS broth were boiled prior to distribution in Hungate tubes (SciQuip, cat. no. 2047-16125), and subsequently flushes with 100% N_2_ with an anaerobic gassing unit (QCAL Messtechnik GmbH, Munich, Germany) for at least 3 min per 10 mL. Both liquid and solid media contained 0.02% (w/v) 1,4-dithiothreitol (Merck, cat. No. DTT-RO) and 0.05% (w/v) L-cysteine (Merck, cat. no. 168149) as reducing agents and 1 mg/L resazurin as oxygen indicator. All media were autoclaved (121°C for 20 min). Anaerobic handling of cultures was performed in an anaerobic chamber (Model AALC, Coy Laboratory Products, Grass Lake, Michigan, USA) containing ∼93% (v/v) N_2_, ∼2% H_2_, ∼5% CO_2_ at room temperature (RT), and agar plates were incubated in an anaerobic jar (Thermo Scientific, cat. no. HP0011A,) along with an anaerobic sachet (Thermo Scientific, AnaeroGen™ cat. no. AN0035A). Incubation of tubes as well as plates was performed at 37°C. For preparing the probiotic solutions, a single bacterial colony was inoculated to the growth medium and incubated until the stationary phase was reached after 24 hours for LGG or 72 hours for AKM. This was followed by a 2% (v/v) culture incubation until the exponential phase was reached after 12 hours for LGG or 48 hours for AKM. The bacterial concentrations were measured with an optical density at 600 nm (OD_600_) with Genesys™ 30 Visible spectrophotometer (Thermo Scientific, cat. no. 840-277000, Waltham, Massachusetts, USA) mounted with a test-tube holder (VWR, cat. no. 634-0911). To ensure high bacterial loads in the probiotic inocula, the bacterial cultures were upconcentrated 40x by centrifugation at 4450 x g for 30 min at RT under anaerobic conditions and resuspended in anaerobic Intralipid®. Intralipid® was used to protect the viable bacterial cells against the acidic environment in the mouse upper gastrointestinal tract [56] and an oil-water emulsion solution was made by mixing the resuspended bacterial cultures with a 3-way stopcock (Braun, Discofix® cat. no. 409511). Pure Intralipid® was used as the probiotic sham (Pro-sham). Small single-use vials of the probiotic solutions were prepared for each mouse to minimize the introduction of oxygen when administering the probiotics. The probiotic solutions were freshly prepared for both the 1^st^ and 2^nd^ inoculation which explain the variances in the bacterial colony forming units (CFU)/mL. Phase contrast microscopy images were taken to check for contamination on the cell morphology level (Figure S11). The total CFU transferred to each mouse at 1^st^ inoculation was LGG: 2.8 × 10^8^ CFU and AKM: 3.8 × 10^8^ CFU and at 2^nd^ inoculation LGG: 5.5 × 10^9^ CFU and AKM: 2.0 × 10^9^ CFU.

### Preparation of donor virome

Fecal viromes were extracted from intestinal content from mice (low-fat diet fed male C57BL/6NCrl and C57BL/6NRj mice) that previously [22] was found with a relative abundance of *A. muciniphila* above 6% and to exhibit inter-vendor variance in their GM profiles [22,57]. The titer of the applied FVT virome was approximately 5.4 × 10^9^ virus-like particles (VLP)/mL (Figure S12 and Table S1) for both 1^st^ and 2^nd^ inoculation and was evaluated by epifluorescence microscopy stained by SYBR™ Gold (Thermo Scientific, cat. no. S11494) as previously described (http://dx.doi.org/10.17504/protocols.io.bx6cpraw). The total VLPs transferred to each mouse per inoculation was 8.0 × 10^8^ VLPs. SM buffer (NaCl 200 mM, MgSO_4_·7H_2_O 16 mM, Tris-HCl 100 mM, pH 7.5) was used as viral sham (Saline).

### Animal study design

In total 48 C57BL/6NTac mice at 4 weeks old (Taconic, Lille Skensved, Denmark) were included in the study (representing 24 males and 24 females). They were ear tagged upon arrival and divided into six groups: LGG+FVT, AKM+FVT, Pro-sham+FVT, LGG+Saline, AKM+Saline, and control (Pro-sham+Saline) (Figure 1). The Saline consisted of SM buffer and Pro-sham consisted of Intralipid® (Fresenius Kabi, Intralipid® 200 mg/mL) that were used to suspend the probiotic bacteria. The mice were housed in open transparent cages with a wire lid (1290D Eurostandard Type III, Scanbur A/S, Karlslunde, Denmark) with access to bottled tap water *ad libitum*, and the cages were enriched with bedding, cardboard housing, tunnel, nesting material, felt pad, and biting stem

(respectively, Cat. no. 30983, 31000, 31003, 31008, 31007, 30968 Brogaarden). The mice were fed *ad libitum* chow diet (Altromin 1324, Brogaarden) during the entire 6 weeks of the study. Health monitoring of animals was performed without revealing any pathogens according to FELASA guidelines [58]. Cages were changed weekly. The mice were housed in male-female pairs (in total 24 cages) to evaluate the effect of sex of the animals on the interventions, increase animal welfare by eliminating aggression between co-housed males, and by consequence also allowed natural mating behavior. After a week of acclimatization, the mice were inoculated orally using a pipette with 50 µl 1M bicarbonate solution (Merck, cat. no. S5761) that 5 min later was followed by oral gavage with 0.15 mL FVT/Saline solutions (FVT/Saline, n = 24). The following day the mice were inoculated orally by gently using a pipette with 100 µL probiotic solution of AKM/LGG/Pro-sham (AKM/Pro-sham, n=16 and LGG/Pro-sham, n=16), which constituted the 1^st^ inoculation. The same procedures were repeated in the 2^nd^ inoculation a week after. The mice were weighted, and fecal samples were taken at several timepoints during the study, amongst other at baseline, 6 days after 2^nd^ intervention, and at termination. The fecal samples were stored at -80°C. One female mouse (representing the AKM+FVT group) was sacrificed following the first probiotic inoculation due to suffering. At termination (9 weeks of age), the remaining 47 mice were anesthetized with a hypnorm/midazolam mixture. Both hypnorm (Hypnorm BN: P736/005, VetaPharma Ltd, Leeds, UK) and midazolam were mixed with sterile water in a ratio of 1:1 (BN: 353 0418, Braun, Melsungen, Germany). The animals were euthanized by cervical dislocation. The mesenteric lymph node (MLN) was sampled in ice cold PBS and 2 cm of the distal ileum was sampled in two pieces and snap frozen in liquid nitrogen and stored in -80°C. Surgical equipment used for tissue and fecal sampling during terminal procedures was sterilized between each animal. Pups, both born and *in utero* were counted and euthanized by decapitation. The study was approved by the Danish Competent Authority, The Animal Experimentation Inspectorate, under the Ministry of Environment and Food of Denmark, and performed under license No. 2017-15-0201-01262 C1-3. Procedures were carried out in accordance with the Directive 2010/63/EU and to the Danish law LBK Nr 726 af 09/091993, and housing conditions as earlier described [22].

### Gene expression assay

Ileum (1 cm) pieces were transferred to tubes (Mpbio, cat. no. FastPrep® 50-76-200) including 0.6g glass beads (Sigma-Aldrich, cat. no. G4649), 600 µl lysis binding solution concentrate (Invitrogen™, cat. no. AM1830) and 0.7% β-mercaptoethanol (Sigma-Aldrich, cat. no. M6250) and homogenized on the FastPrep-24™ Classic Instrument (Mpbio, Irvine, California, USA) with 4 x (45 sec at speed 6.5 m/s) runs. The homogenate was centrifuged at 16,000 x g and the supernatant was frozen at -20°C for at least 24 hours before purification of RNA using the MagMax™ Express Magnetic Particle Processor (Applied Biosystems™, Waltham, Massachusetts, USA) using manufacturer’s instructions (Invitrogen™, cat. no. AM1830). RNA purity and concentration were assessed using DeNovix DS11 Fx+ Spectrophotomer (DeNovix, Wilmington, USA) and intact 18S rRNA and 28S rRNA bands was visually inspected on a 1.4% agarose gel. cDNA was synthesized from 500 ng total RNA with the High-capacity cDNA reverse Transcription Kit (Applied Biosystems™, Waltham, Massachusetts, USA) in a reaction volume of 20 µl following manufacturers recommendations. 2 -RT controls were prepared without the Reverse transcriptase enzyme and cDNA samples was diluted 8x after synthesis. High throughput qPCR was run on duplicates on the Biomark HD system (Fluidigm Corporation, South San Francisco, California, USA) on 2x 96.96 IFC chips on pre-amplified cDNA duplicates using manufacturer’s instructions with minor adjustments as previously described [59]. The majority of the primers in the Ileum primer panel was previously published [17]. 81 primer assays (74 candidate genes, 7 reference genes, 1 gDNA control assay) (see Supplementary file 1 for the full list) were present with one product and had a sufficient efficiency between 75-110%. In addition, the MVP1 gene assay was included to control for gDNA contamination [60]. qPCR data was analyzed as previously described and normalized to the four most stable reference genes: Sdha, Tuba, Pgk1, Ppia [17,59]. A log2 fold change threshold was set to 0.5 and the FDR p-value to 0.05 for gene expressions to be included in the analysis.

### Cell isolation and flow cytometry (FACS)

Directly after euthanasia of the mice, the mesenteric lymph node (MLN) was placed in ice cold PBS. Single cell suspensions were prepared by disrupting the lymph node between two microscope glasses and passing it through a 70 μm nylon mesh. After washing and resuspension, 1×106 cells were surface stained for 30 min with antibodies for Percp-Cy5.5 conjugated CD11c, PE-conjugated CD86, APC-conjugated CD11b, and FITC-conjugated CD103 (all antibodies were purchased from eBiosciences, San Diego, CA USA) for the detection of tolerogenic dendritic cells (DCs). For the detection of T cell subsets, 1×106 cells were initially surface stained for 30 min with FITC-conjugated CD3, PercP-Cy5.5-conjugated CD4, and APC-conjugated CD8a (ebiosciences), then fixate and permeabilized with the FoxP3/Transcription Facter Staining Buffer Set (ebiosciences), and finally stained for 30 min with PE-conjugated intracellular forkhead box P3 (FOXP3) (ebioscience). Analysis was performed using an Accuri C6 flow cytometer (Accuri Cytometers, Ann Arbor, MI, USA).

### Pre-processing of fecal samples

Fecal samples from three different timepoints were included to investigate GM changes over time: baseline, after intervention, and at termination. This represented in total 142 fecal samples from the C57BL/6NTac mice. Separation of the viruses and bacteria from the fecal samples generated a fecal pellet and fecal supernatant as earlier described [22].

### Quantitative real-time PCR for measuring probiotic density

The bacterial density of AKM and LGG in the fecal samples was estimated using quantitative real-time polymerase chain reaction (qPCR) with species specific primers (AKM_Fwd: 5’-CCT TGC GGT TGG CTT CAG AT-3’ and AKM_Rev: 5’-CAG CAC GTG AAG GTG GGG AC-3’ [61] and LGG_Fwd: 5’-GCC GAT CGT TGA CGT TAG TTG G-3’ and LGG_Rev: 5’-CAG CGG TTA TGC GAT GCG AAT-3’ [62]) purchased from Integrated DNA Technologies (IDT, Iowa, USA). Standard curves (Table S2) were based on a dilution series of total DNA extracted from monocultures of AKM and LGG. The qPCR results were obtained using the CFX96 Touch Real-Time PCR Detections System (Bio-Rad Laboratories, Hercules, California, USA) and the reagent SsoFast™ EvaGreen® Supermix with Low ROX (Bio-Rad Laboratories, cat. no. 1725211), and run as previously described [63].

### Bacterial DNA extraction, sequencing and pre-processing of raw data

The Bead-Beat Micro AX Gravity kit (A&A Biotechnology, cat. no. 106-100 mod.1) was used to extract bacterial DNA from the fecal pellet by following the instructions of the manufacturer. The final purified DNA was stored at -80ºC and the DNA concentration was determined using Qubit HS Assay Kit on the Qubit 4 Fluorometric Quantification device (Invitrogen, Carlsbad, California, USA). The bacterial community composition was determined by Illumina NextSeq-based high-throughput sequencing (HTS) of the 16S rRNA gene V3-region, as previously described [22]. Quality-control of reads, de-replicating, purging from chimeric reads and constructing zOTU was conducted with the UNOISE pipeline [64] and taxonomically assigned with Sintax [65] (not yet peer reviewed). Taxonomical assignments were obtained using the EZtaxon for 16S rRNA gene database [66]. Code describing this pipeline can be accessed in github.com/jcame/Fastq_2_zOTUtable. The average sequencing depth after quality control (Accession: PRJEB52388, available at ENA) for the fecal 16S rRNA gene amplicons was 47,526 reads (min. 6,588 reads and max. 123,389 reads).

### Viral DNA extraction, sequencing and pre-processing of raw data

The sterile filtered fecal supernatant was concentrated using Centriprep® filter units (Merck, cat. no. 4311 and cat. no. 4307). This constituted the concentrated virome. Due to a permanent stop in the production of Centriprep® filter units at the manufacturer, we were forced to use residual stocks of filter size 30 kDa (cat. no. 4307) for 42% of the samples (Table S3). The samples were all centrifuged at 1500 x g at 15ºC until approx. 500 µL concentrated virome sample was left, the filter was removed with a sterile scalpel and stored along with the concentrated virome at 4ºC. Viral DNA was extracted, multiple displacement amplification (MDA, to include ssDNA viruses), and Illumina NextSeq sequencing data generated as previously described [22]. The average sequencing depth after quality control (Accession: PRJEB52388, available at ENA) for the fecal viral metagenome was 209,641 reads (min. 21,580 reads and max. 510,332 reads. The raw reads were trimmed from adaptors and the high quality sequences (>95% quality) using Trimmomatic v0.35 [67] with a minimum size of 50nt were retained for further analysis. High quality reads were de-replicated and checked for the presence of PhiX control using BBMap (bbduk.sh) (https://www.osti.gov/servlets/purl/1241166). Virus-like particle-derived DNA sequences were subjected to within-sample de-novo assembly-only using Spades v3.13.1 [68] and the contigs with a minimum length of 2,200 nt, were retained. Contigs generated from all samples were pooled and de-replicated at 90% identity using BBMap (dedupe.sh). Prediction of viral contigs/genomes was carried out using VirSorter2 [69] (“full” categories | dsDNAphage, ssDNA, RNA, Lavidaviridae, NCLDV | viralquality ≥ 0.66), vibrant [70] (High-quality | Complete), and checkv [71] (High-quality | Complete). Taxonomy was inferred by blasting viral ORF against viral orthologous groups (https://vogdb.org) and the Lowest Common Ancestor (LCA) for every contig was estimated based on a minimum e-value of 10e^-5^. Phage-host prediction was determined by blasting (85% identity) CRISPR spacers and tRNAs predicted from >150,000 gut species-level genome bins (SGBs) [72,73] ([73], not yet peer reviewed). Following assembly, quality control, and annotations, reads from all samples were mapped against the viral (high-quality) contigs (vOTUs) using the bowtie2 [74] and a contingency-table of reads per Kbp of contig sequence per million reads sample (RPKM) was generated, here defined as vOTU-table (viral contigs). Code describing this pipeline can be accessed in github.com/jcame/virome_analysis-FOOD.

### Bioinformatic analysis of bacterial and viral DNA sequences

Initially the RPKM normalized dataset was purged for viral contigs which were detected in less than 5% of the samples, but the resulting dataset still maintained 99.5% of the total reads. Cumulative sum scaling (CSS) [75] was applied for the analysis of β-diversity to counteract that a few viral contigs represented a majority of count values, since CSS have been benchmarked with a high accuracy for the applied metrics [76]. CSS normalization was performed using the R software using the metagenomeSeq package [77]. β-diversity analysis was based on raw read counts and statistics were based on ANOVA. R version 4.01 [78] was used for subsequent analysis and presentation of data. The data are uploaded as supplementary data (www.osf.oi/tm2a5). The main packages used were phyloseq [79], vegan [80], deseq2 [81], ampvis2 [82] (not yet peer reviewed), ggpubr [83], mctoolsr (https://github.com/leffj/mctoolsr/), and ggplot2 [84]. B-diversity was represented by Bray Curtis dissimilarity and statistics were based on PERMANOVA. A linear model (y∼FVT+probiotics+sex), similar to ANOVA, was applied to assess the statistically differences between the treatment groups of gene expression, bacterial abundance, immune cell counts, and fertility rate. Two treatment groups were applied in the model; a FVT group (levels: control and FVT) and a probiotic group (levels: control, AKM and LGG). The sex of the animal was added as an additional factor, except for fertility outcome analysis where only females were included. For binary outcomes a generalized logistic regression model was applied.

## Acknowledgements

We thank the animal caretakers Helene Farlov and Mette Nelander at Section of Experimental Animal Models (University of Copenhagen, Denmark) for taking care of the animals during the study and assisting with the animal handling.

## Author contributions

TSR, CMJM, AKH, and DSN conceived the research idea and designed the study; TSR, CMJM, MRD, LSFZ, performed the experiments TSR, CMJM, MRD, RRJ, LSFZ, JLCM, LHH, AKH, and DSN performed laboratory and data analysis; TSR wrote the first draft of the manuscript. All authors critically revised and approved the final version of the manuscript.

## Data availability statement

Supplementary materials and raw data used for the analysis can be accessed through doi: 10.17605/OSF.IO/TM2A5 (www.osf.oi/tm2a5). Raw sequencing data can be accesses at ENA with project ID: PRJEB52388 (https://www.ebi.ac.uk/ena/browser/).

