## Supplemental figures for "Fecal virome transfer improves proliferation of commensal gut *Akkermansia muciniphila* and unexpectedly enhances the fertility rate in laboratory mice"

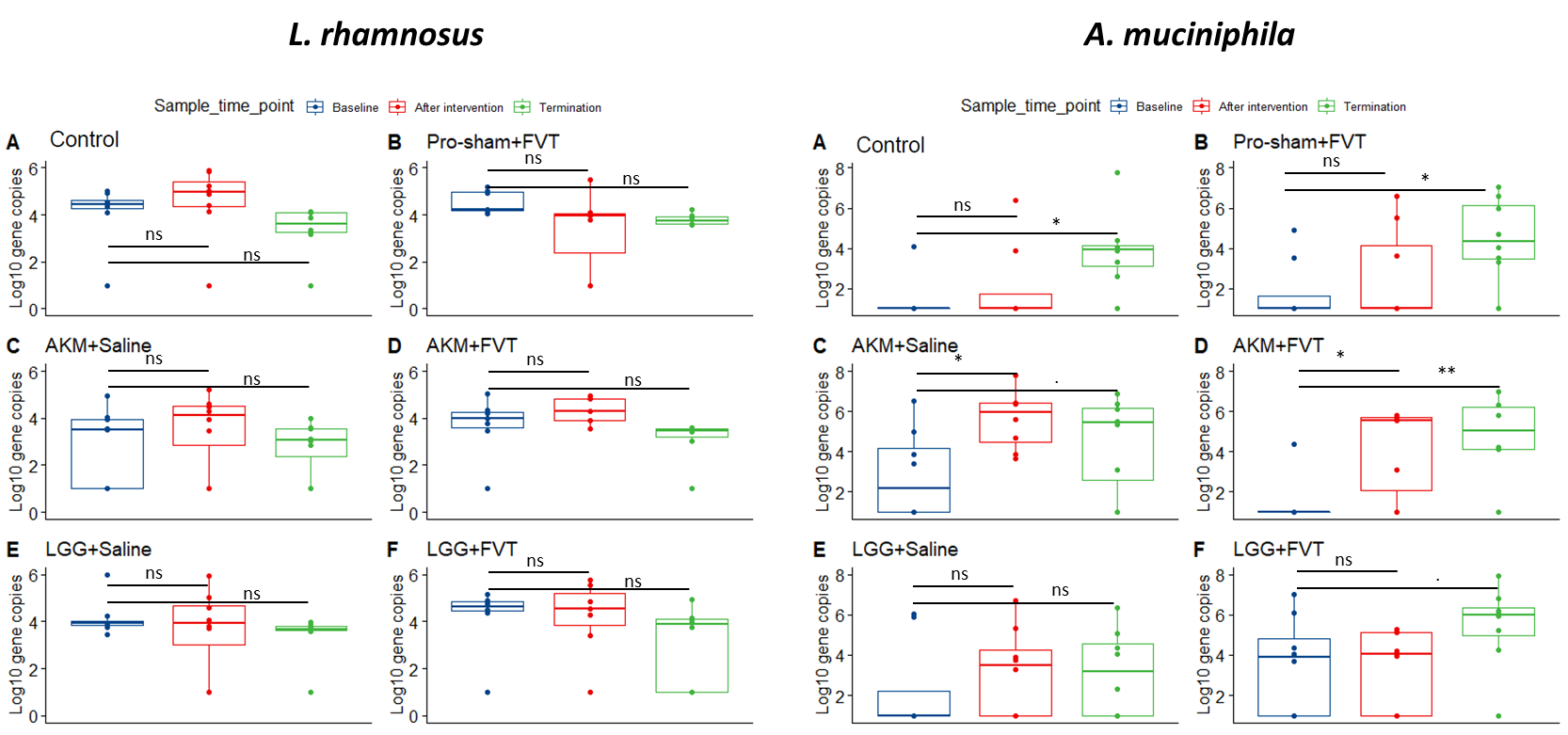

Figure S1: qPCR with L. rhamnosus and A. muciniphila specific primers were used to assess the abundance of gene copies pr. gram feces over the time span of baseline, after intervention, and termination. Here for each group showing the development of gene copies over time. Abbreviations: “ns” = not significant, “.” = p < 0.1, “*” = p < 0.05, “**” = p < 0.001.

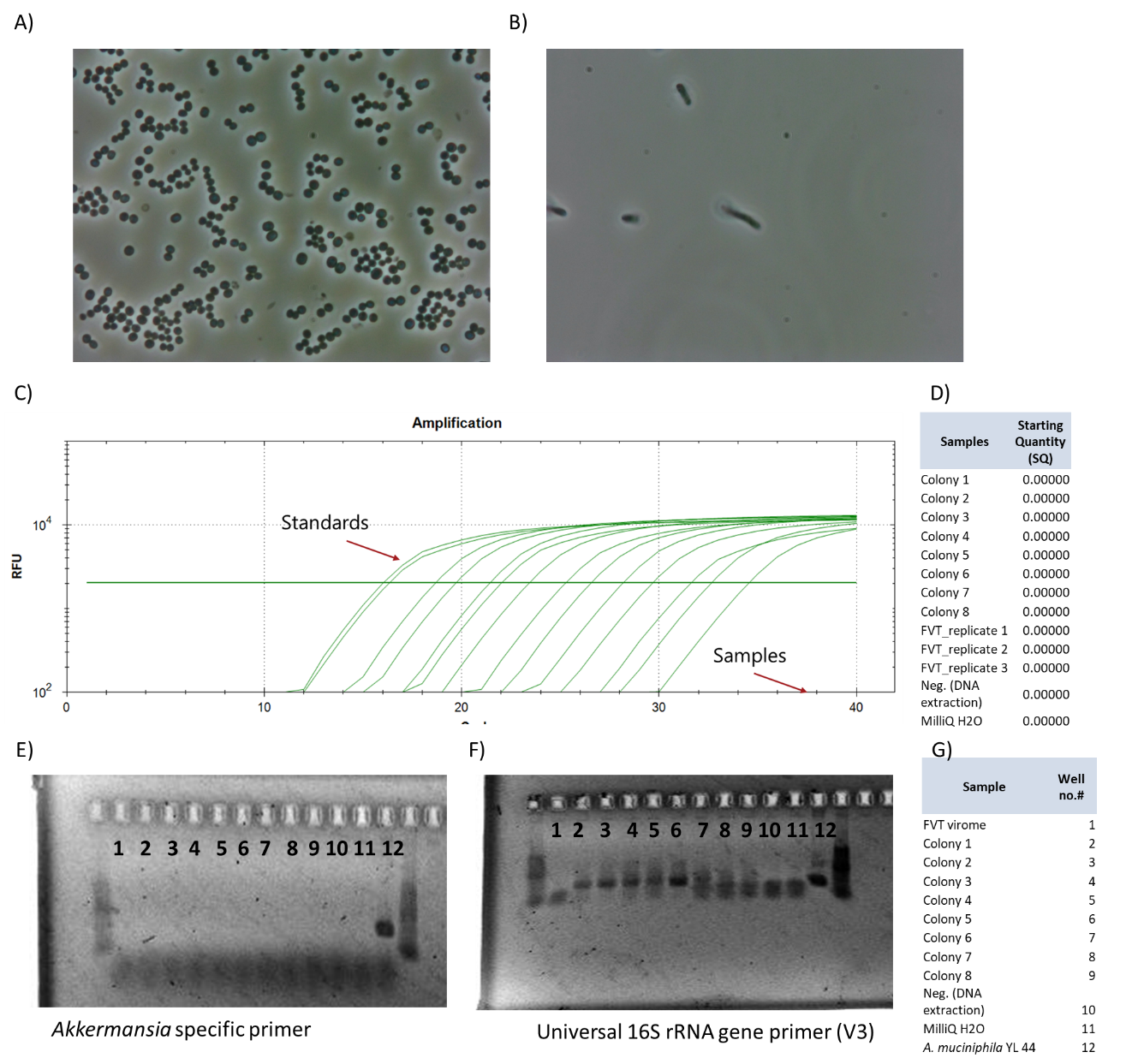

Figure S2: A) and B) Phase contrast microscopy images representing two different cell morphologies detected in the eight bacterial colonies found after incubation of FVT virome on GAM plates. C) Showing the standard curve and samples D) readouts of qPCR using Akkermansia specific primers, where no traces of Akkermansia in the FVT donor virome, nor the isolated colonies were observed. E) PCR products on an agarose gel after running PCR with Akkermansia specific primers as well as universal 16S rRNA gene primers (V3) F) of samples listed in G). Neither the PCR detected any traces of Akkermansia in the FVT virome, nor the isolated colonies.

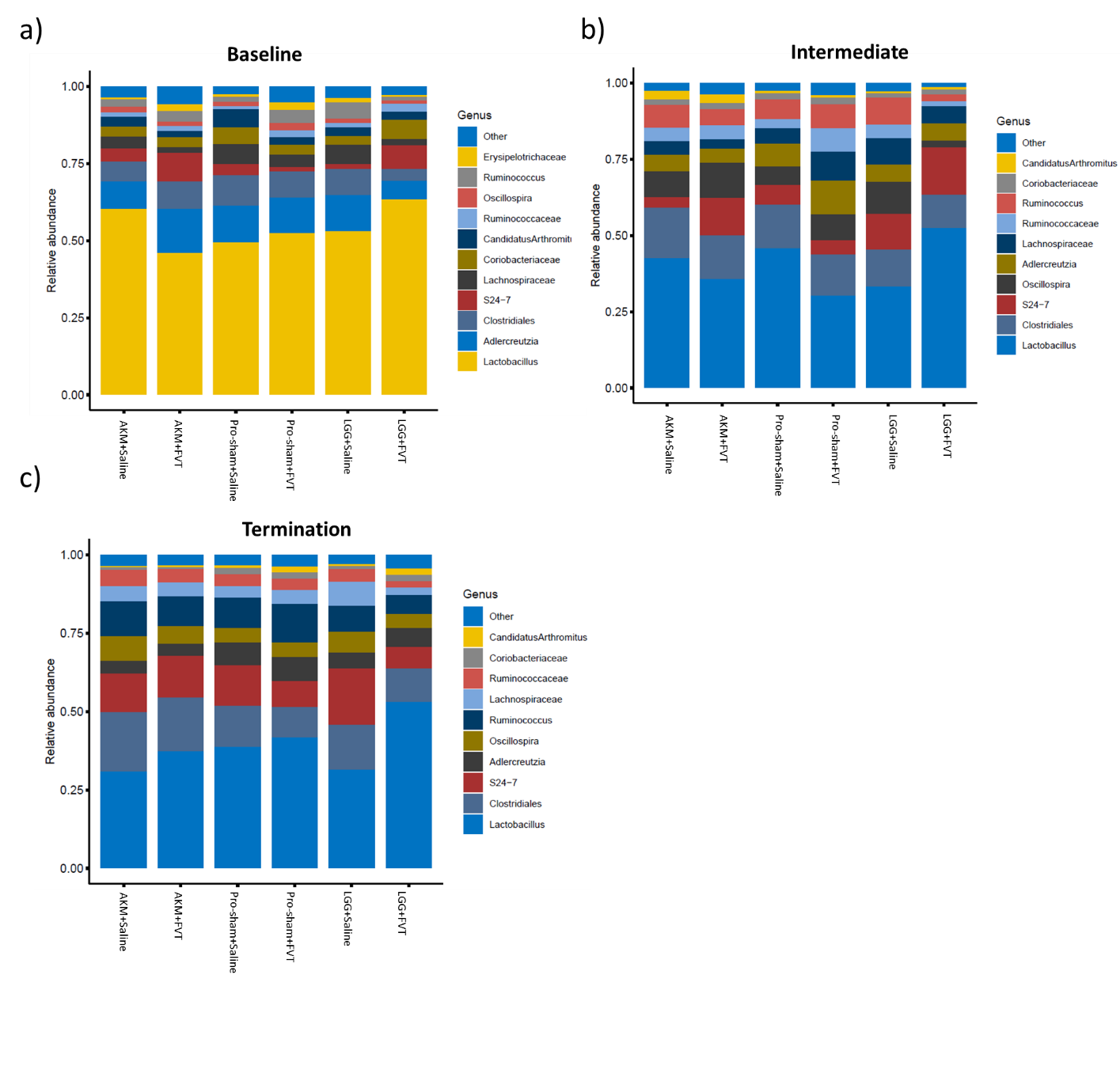

Figure S3: The relative abundance of the most abundant bacterial taxa in the different treatment groups of the time span of baseline, after intervention, termination. Abbreviations: Lacticaseibacillus rhamnosus GG = LGG, Akkermansia muciniphila = AKM, fecal virome transplantation = FVT, Pro-sham = probiotic sham, Saline = SM buffer.

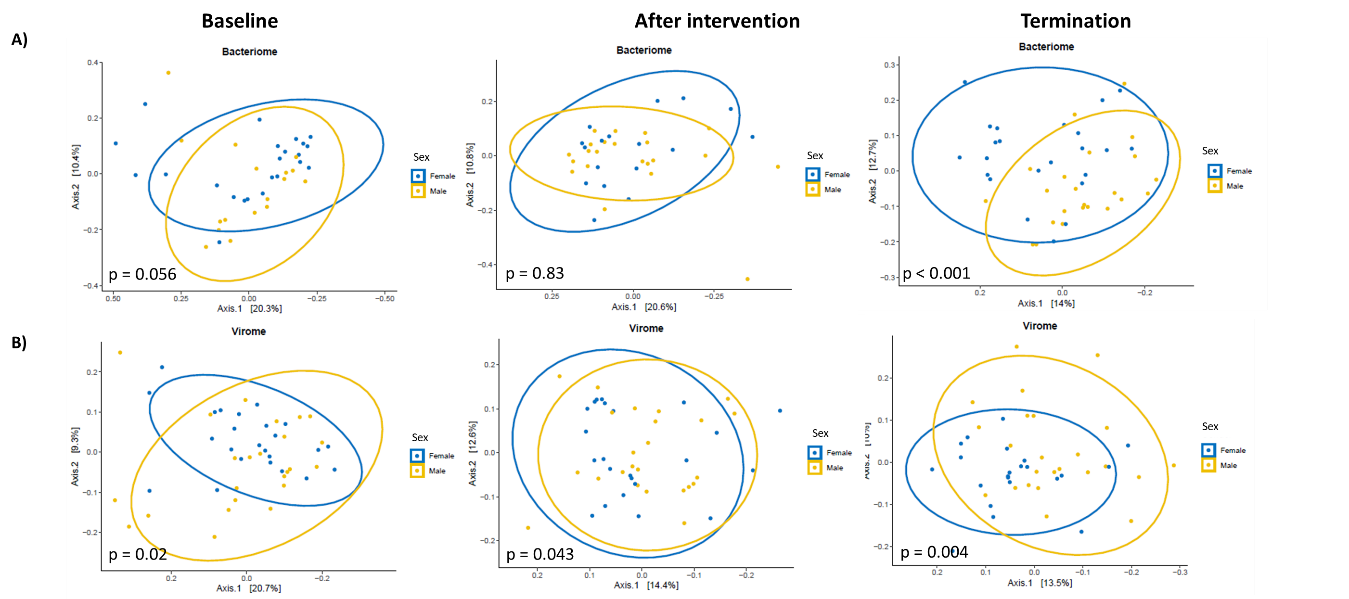

Figure S4: Beta-diversity PCoA plots based on Bray Curtis dissimilarity of both A) Bacteriome and B) Virome at all three time points showing the effect of the sex of the animals at baseline, after intervention, and termination.

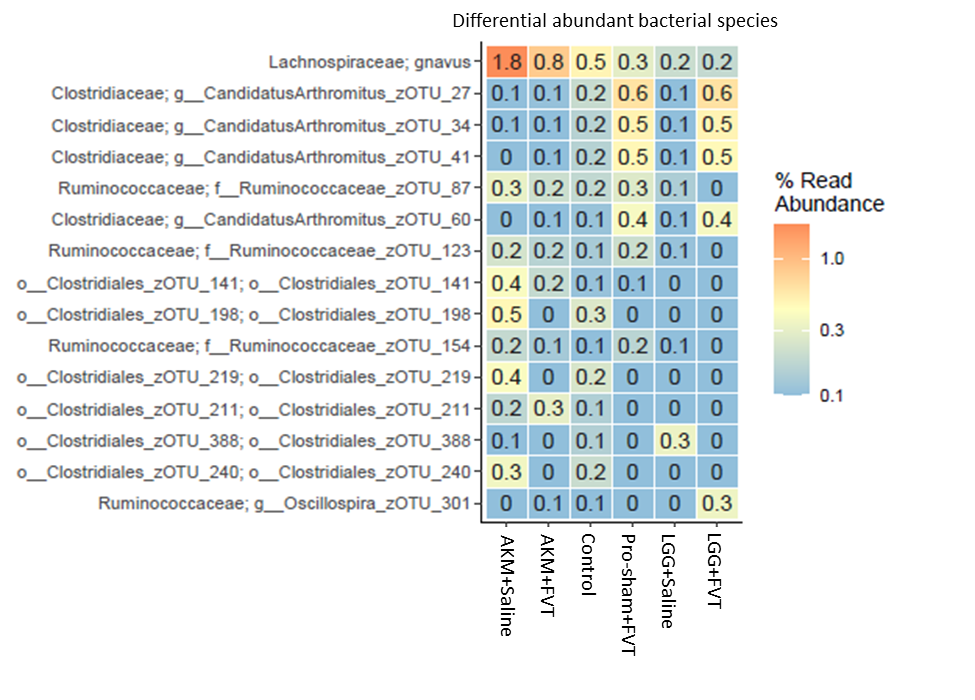

Figure S5: Differential abundance analysis of the bacteriome data comparing the different treatment groups. R. gnavus appeared significantly (p < 0.05) higher in relative abundance compared in the mice treated with AKM and/or FVT.

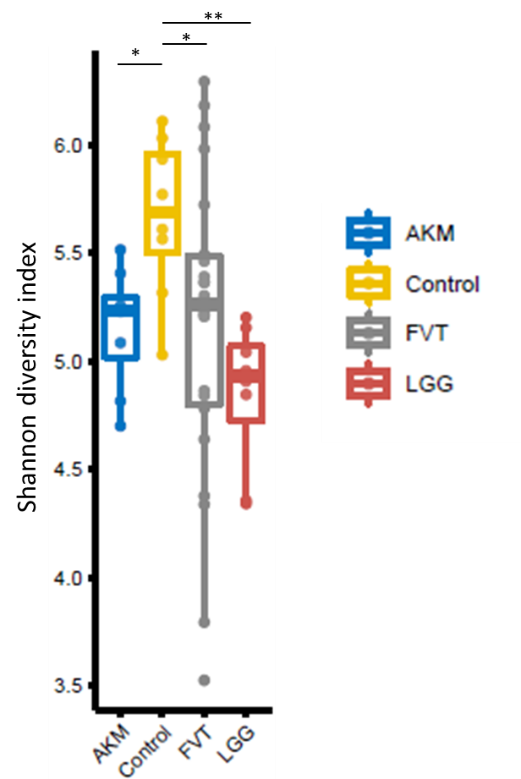

Figure S6: Alpha-diversity (viral diversity) plot based on the Shannon diversity index. Mice that received FVT are here merged to one group and compared with mice that received AKM or LGG as well as the control (Pro-sham+Saline) mice. “*” = p < 0.05, “**” = p < 0.007.

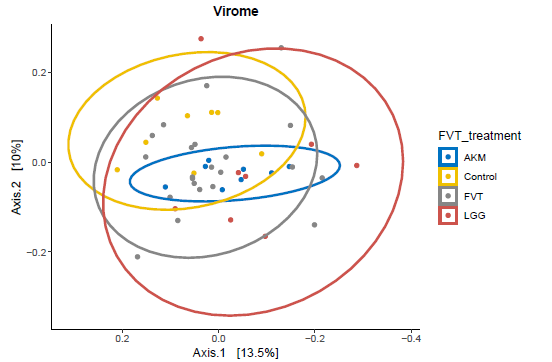

Figure S7: Beta-diversity (viral composition) PCoA plots based on Bray Curtis dissimilarity showing that AKM (p = 0.025) and LGG (p = 0.014) at termination significantly affected the viral composition.

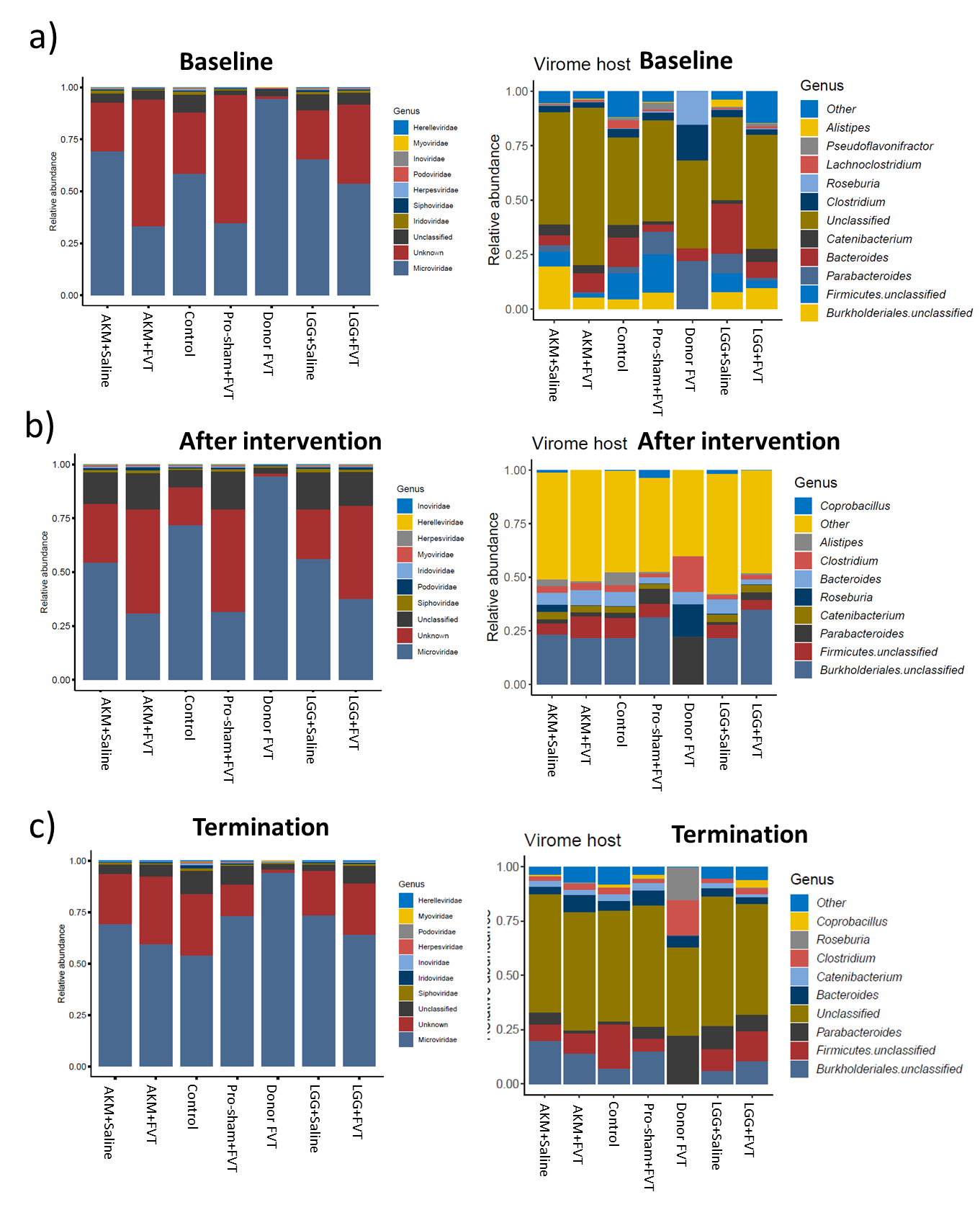

Figure S8: The relative abundance of the most abundant viral taxa and host-predicted taxa in the different treatment groups of the time span of a) Baseline, b) after intervention), and c) termination. Abbreviations: Lacticaseibacillus rhamnosus GG = LGG, Akkermansia muciniphila = AKM, fecal virome transplantation = FVT, Pro-sham = probiotic sham, Saline = SM buffer

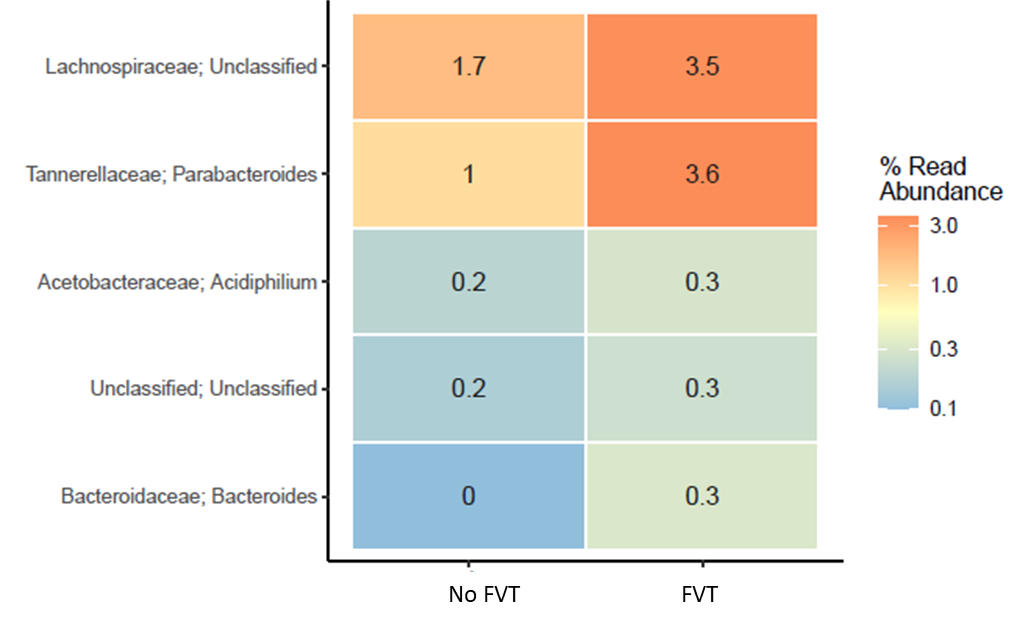

Figure S9: Differential abundance analysis of the predicted hosts based on the metavirome data, which here is comparing the mice treated with FVT with the mice receiving no FVT. The FVT-treated mice had a significant (p < 0.05) increase in viruses that was predicted to infect *Lachnospiraceae*, *Bacteroides*, and *Parabacteroides*.

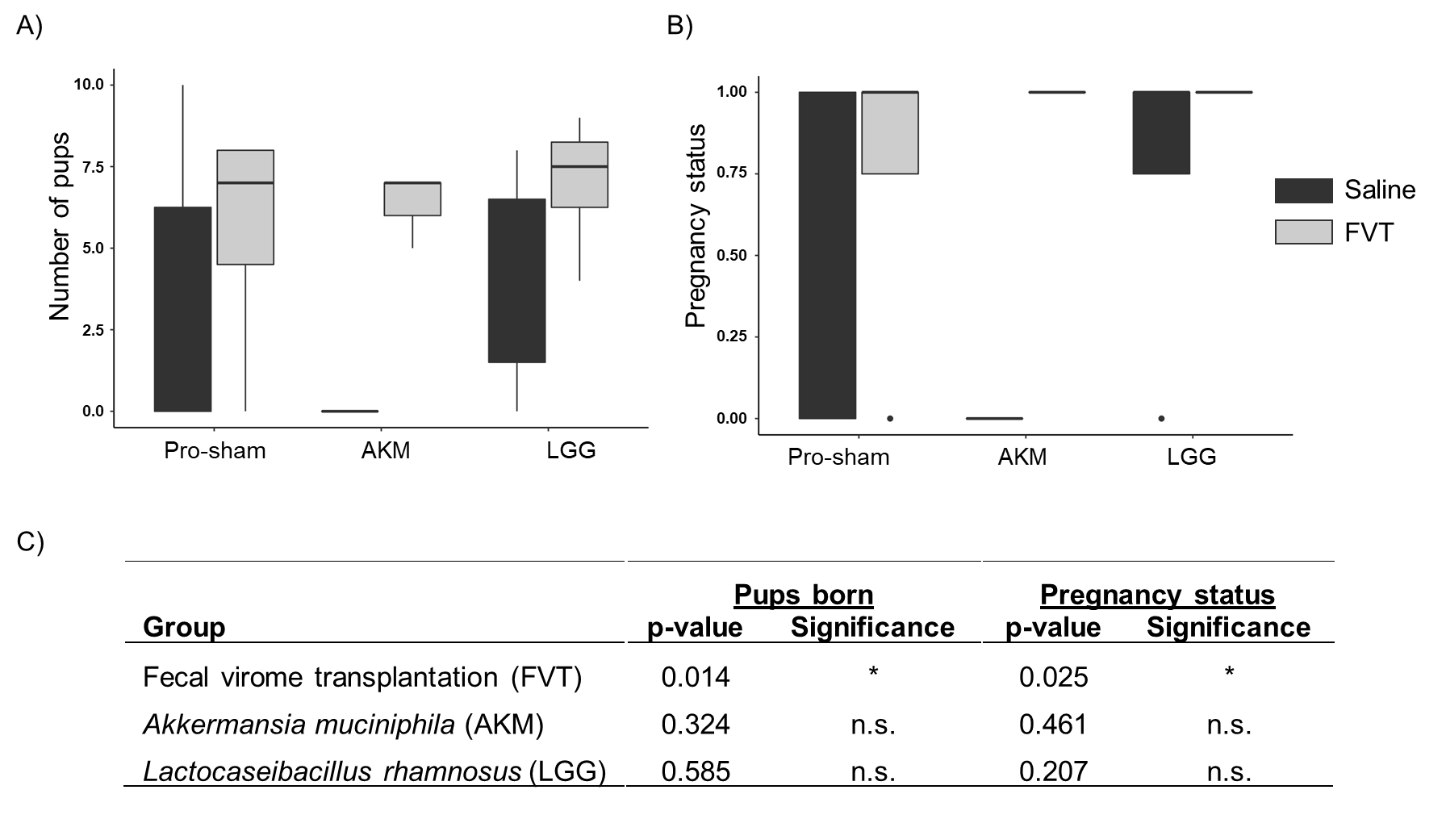
Figure S10: Bar plots of the fertility rate and pregnancy status. A) The observed number of pups (born or as fetuses) based on a linear model (y~FVT+probiotics). B) The distribution of the binary event of pregnancy (pregnant or not). Only the 23 female mice that received either FVT/Saline along with probiotic solutions of AKM/LGG/Pro-sham was included in the C) statistical analysis that was based on a generalized logistic regression model. Abbreviations: Lacticaseibacillus rhamnosus GG = LGG, Akkermansia muciniphila = AKM, fecal virome transplantation = FVT, Pro-sham = probiotic sham.

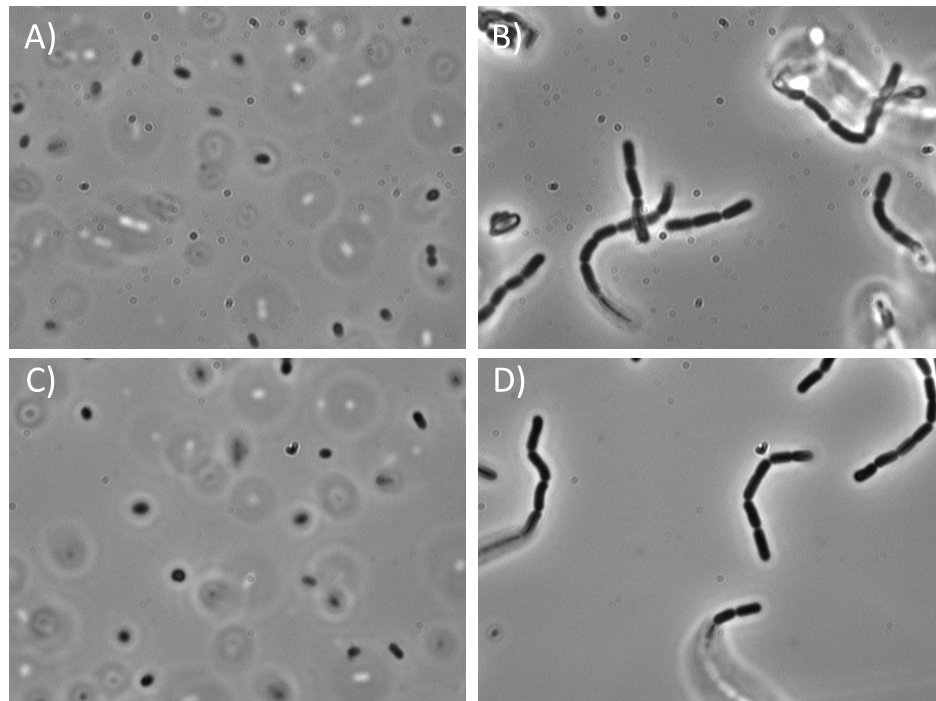

Figure S11: Phase contrast microscopy (100x magnification) images of probiotic cultures of A. muciniphila YL-44 (A and C) and L. rhamnosus GG (B and D) for respectively 1^st^ and 2^nd^ inoculation.

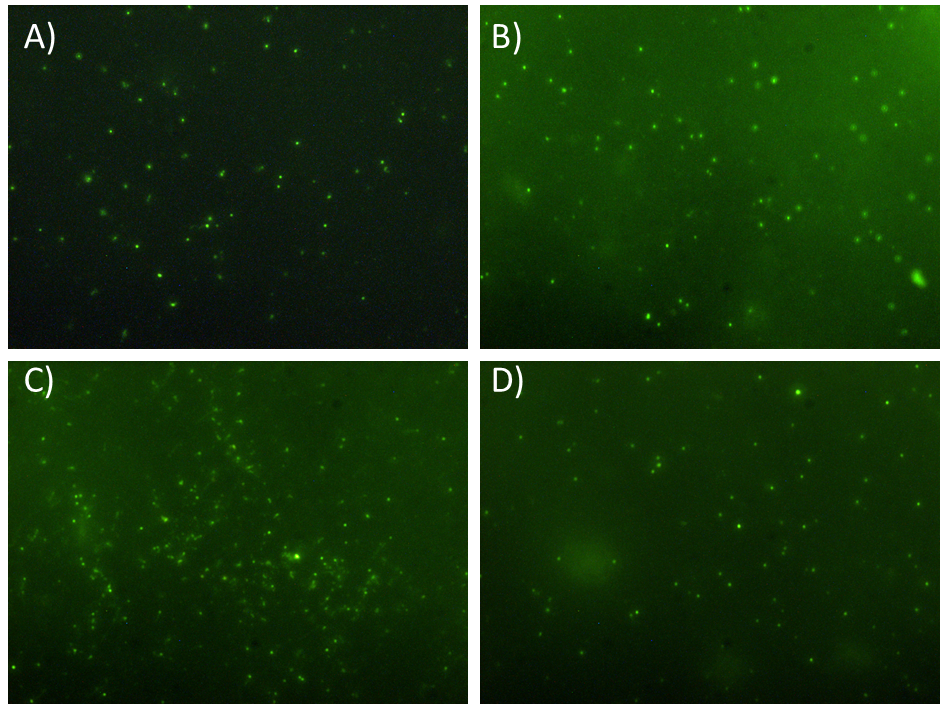

Figure S12: Epifluorescence microscopy images of the applied FVT virome that was stained with SYBR Gold to count VLP/mL, as calculated in Table S1. Image ID according to TableS1 19, 20, 22, 26 at respectively A), B), C) and D).
