## Supplementary material for "Fecal virome transfer improves proliferation of commensal gut *Akkermansia muciniphila* and unexpectedly enhances the fertility rate in laboratory mice": Supplemetary file 1

| Gene name | FW | RV | Amplicon length | Gene function /pathway |
|---|---|---|---|---|
| *Arg1* | ATGGGCAACCTGTGTCCTTT | TCTACGTCTCGCAAGCCAAT | 127 | M2 Macrophages |
| *Ccl2* | CAGCTCTCTCTTCCTCCACC | TGGGATCATCTTGCTGGTGA | 155 | MCP-1 Monocyte attractant |
| *Ccl3* | ACCATGACACTCTGCAACCA | CAACGATGAATTGGCGTGGA | 106 | MIp1-a Cytokine (lukocyte attractant |
| *Ccr10* | AAACCCTTGTAGCCAGAGATGG | CGTACCCGGAGTAAAGTCCC | 70 | Treg skin homing marker. Potentially gut |
| *Ccr9* | CTCTGGTCTGCCTTTGCCAC | GGCATCATGGTGGATCGGAC | 84 | T cell gut homing marker |
| *Cd38* | ACTGGAGAGCCTACCACGAA | AGTGGGGCGTAGTCTTCTCT | 179 | M1 macrophages |
| *Cd3e* | CCTCCTAGCTGTTGGCACTT | GGGCACGTCAACTCTACACT | 90 | T cells |
| *Cd8a* | GGATTGGACTTCGCCTGTGA | TGGGACATTTGCAAACACGC | 130 | CD8 T cells |
| *Cdh1* | GAGACCAGTTTCCTCGTCCG | AGCAGCTCTGGGTTGGATTC | 137 | Cadhedrin-1 /E-cadhedrin Instestinal barrier function |
| *Cgn* | TGAATCCGGGAGCACTGATCT | ACGAAGGGTGCCCATTTCAG | 151 | gut barrier |
| *Cldn1* | TCGACTCCTTGCTGAATCTGA | CAGCCATCCACATCTTCTGC | 159 | Gut barrier |
| *Cldn2* | CGCCTTTCTCTGGACCTAGT | CTTGCTTCTTGGATCCGAGC | 192 | Gut barrier |
| *Cldn7* | GCCTTGGTAGCATGTTCCTG | TTTGCTTTCACTGCCTGGAC | 180 | Gut barrier |
| *Ctla4* | ATGGCTTGTCTTGGACTCCG | ACCACTGAAGGTTGGGTCAC | 137 | Tregs |
| *Ctnnb1* | GAGCACATCAGGACACCCAA | CCGAGCAAGGATGTGGAGAG | 122 | Beta-catenin, gut barrier |
| *Cxcl1* | TGCACCCAAACCGAAGTCAT | CTCCGTTACTTGGGGACACC | 122 | Neutrophil granolucytes |
| *Cxcl16* | CCCAGATACCGCAGGGTACTT | TTCCCATGACCAGTTCCACA | 181 | NK/iNKT |
| *Cxcl10* | AAGTGCTGCCGTCATTTTCT | CCTATGGCCCTCATTCTCAC | 129 | Cytokine |
| *Cxcr6* | TGGAACAAAGCTACTGGGCT | TCGTAGTGCCCATCGTACAG | 81 | NK/iNKT |
| *Epx* | ACCGAACCATAACAGGGAGATG | CTGATTGGAGACATCCCGGAC | 182 | Eosinophil granulocytes |
| *Fasl* | AAGGAACTGGCAGAACTCCG | ACTCCAGAGATCAGAGCGGT | 151 | Cytotocox CD8 Tcells |
| *Ffar1* | CTGGGCATCAACATACCCGT | AGCAGAAGGCAGTGATGACC | 133 | SCFR |
| *Ffar2* | AGGGAGGAATCACAGGAAACG | CTTGGGCAAGTTCAGGGGTT | 165 | Short chain fatty acid signalling |
| *Ffar3* | CAGAGTGCCAGTTGTCCAAT | GCAAAAGTAAGTCCACAGCCA | 188 | Free fatty acids receptor |
| *Ffar4* | TGCCCCTCTGCATCTTGTTC | GGTTGGGCCAATCCAATGTG | 90 | SCFR |
| *Fgf15* | ACGGCAAGATATACGGGCTG | GGCTTGGCCTGGATGAAGAT | 133 | GM signalling |
| *Foxp3* | AGAGAGAAGTGGTGCAGTCTC | GAGTACTGGTGGCTACGATG | 159 | Tregs;CD4 |
| *Fxr* | GCTGAGACTGGGTACCAGGG | TCGGAAGAAACCTTTGCAGCC | 183 | GM signalling |
| *Gapdh* | GGAGAGTGTTTCCTCGTCCC | ATGAAGGGGTCGTTGATGGC | 136 | reference gene |
| *Gata3* | GCTACGGTGCAGAGGTATCC | CAGAGATCCGTGCAGCAGAG | 75 | ILC;CD4; Th2 |
| *Gcg* | TCTACACCTGTTCGCAGCTC | GTCCTCATGCGCTTCTGTCT | 172 | Insulin signalling |
| *Gzmb* | AGAGGGGGTACAAGGTCACA | CATGTCCCCCGATGATCTCC | 187 | CD8 Tcells |
| *Hprt* | TCAGTCAACGGGGGACATAAA | GGGGCTGTACTGCTTAACCAG | 122 | ref gene |
| *Icam1* | CTGTGCTTTGAGAACTGTGGC | CAGGGTGAGGTCCTTGCCTA | 129 | Cellular adhesion - immune cells |
| *Ifng* | TTTGAGGTCAACAACCCACAG | GCTTCCTGAGGCTGGATTC | 94 | Innate; NK/iNKT ; Neutrophil granolucytes |
| *Il10* | AGGCGCTGTCATCGATTTCT | ATGGCCTTGTAGACACCTTGG | 104 | Tregs |
| *Il12b* | TTGTTCGAATCCAGCGCAAG | TTCTCTACGAGGAACGCACC | 83 | p40 subunit |
| *Il15* | ACAGCTCAGAGAGAATCCACC | ATGAGCTGGCTATGGCGATG | 187 | CD8 Tcells |
| *Il17a* | TGAGTCCAGGGAGAGCTTCA | CGCTGCTGCCTTCACTGTA | 80 | Th17 |
| *Il1b* | GCAACTGTTCCTGAACTCAACT | ATCTTTTGGGGTCCGTCAACT | 89 | Th1 |
| *Il2* | GAAACTCCCCAGGATGCTCA | CGCAGAGGTCCAAGTTCATCT | 99 | T cell differentiation |
| *Il27* | GTCCACAGCTTTGCTGAATCT | CGAAGTGTGGTAGCGAGGAA | 149 | il30 |
| *Il33* | GGGCTCACTGCAGGAAAGTA | TTTGCCGGGGAAATCTTGGA | 115 | Th2 |
| *Il4* | CCTGGATTCATCGATAAGCTG | TCCATTTGCATGATGCTCTT | 93 | M2 Macrophages; Th2 |
| *Il6* | GACAAAGCCAGAGTCCTTCAGA | AGGAGAGCATTGGAAATTGGGG | 113 | Th2 |
| *Irf3* | CCACAAGGACAAGGACGGAG | CCACATTTCCCCCATGCAGA | 124 | gut cell signalling |
| *Itgax* | GAGCCAGAACTTCCCAACTG | ACCCGAGCCATCAATCAGG | 79 | Dendritic cells |
| *Muc1* | AGTACCAAGCGTAGCCCCTA | GTGGGGTGACTTGCTCCTAC | 118 | Mucin layer |
| *Muc2* | TATGCCAGGCCAGGAGTTTA | GCAAGGCAGGTCTTTACACA | 82 | Mucin layer |
| *Muc4* | GTCCACTTCTTCCCCATCTCG | CCATTGTGACAGTAGCCCTCA | 173 | Mucin layer |
| MVP1 | GGAGCCCAGTGTAGAAGAGCA | AGCCAGCGAACCATATCCTGA | 87 | MVPA1 from Laurell et all 2012 nucleic acid res |
| *Myd88* | CCAGGTGTCCAACAGAAGC | CTTGGTGCAAGGGTTGGTAT | 114 | Tlr signalling |
| *Nfkb* | GGCAGGTATTTGACATACTAAATG | TGCAGAGTTGTAGCCTCGTG | 117 | transcription factor innate and adaptive immunity |
| *Nfkbia* | GAGCGAGGATGAGGAGAGCTA | GGCCTCCAAACACACAGTCA | 83 | IκBα (regulates NFkb) |
| *Nod2* | TGGCCCTACAGCTGGATTAC | TTGTTGTTGAAGAGACTGGCTA | 187 | CARD15 - pattern recognition receptor |
| *Nos2* | GTGACCATGGAGCATCCCAA | TCGAACTCCAATCTCGGTGC | 159 | M1 macrophages |
| *Ocld* | GCTGCTGCTGATGAATATAATAGA | TCCCACCATCCTCTTGATGT | 120 | Gut barrier |
| *Pgk1* | GGTGTTGCCAAAATGTCGCT | GGACTTGGCTCCATTGTCCA | 183 | ref gene |
| *Pla2g2a* | GGGGCCAAATCACCTGTTCT | GTTCCGGGCGAAACATTCAG | 92 | antimicrobial factor |
| *Ppara* | AACATCGAGTGTCGAATATGTGG | CCGAATAGTTCGCCGAAAGAA | 99 | immune and lipid metabolism |
| *Pparg* | TTCAGAAGTGCCTTGCTGTG | CCAACAGCTTCTCCTTCTCG | 84 | immune and glucose metabolism |
| *Ppia* | CCACCGTGTTCTTCGACATC | AGTGCTCAGAGCTCGAAAGT | 113 | reference gene |
| *Prkaa2* | GCAAAGTGAAGACTACCAGGTG | GTAATCCACGGCAGACAGGA | 163 | AMPK catalytic subunit. Lipid and insulin metabolism |
| *Prf1* | ACACAGTAGAGTGTCGCATGT | GCCGTGATAAAGTGCGTGC | 163 | NK and cytTcells - induce apoptosis in infected cells |
| *Pyy* | GCTTCTCCCACCTTCCATCT | AGACAGGCGAGCAGGATTAG | 121 | Gasttic peptide |
| *Reg3a* | TGGGCTCCATGATCCAACAA | CTGTCAGACTCCCACAGTGAC | 140 | antimicrobial factor |
| *Retnlb* | CTGTCCTGCTGGGATGGT | CCAGTCCATGACTGAGCACT | 109 | Gut barrier |
| *Rorc* | TACCCTACTGAGGAGGACAGG | AATGGGGCAGTTCTGCTGAC | 200 | CD4 Tcells |
| *Sdha* | ATTGCTACTGGGGGCTACGG | GTCCTGGCAAGGCAAACCAG | 108 | ref gene |
| *Stat4* | GAAGTACCTCTACCCTGACATTCC | AGGGGACGTAACCCTTGTCT | 113 | Th1 |
| *Stat5* | GGTCCCTGAGTTCGTCAATG | GGTTGGGTGGGTACATGTTG | 116 | Immunity transcription factor |
| *Tbp* | ACCTTATGCTCAGGGCTTGG | TGCCGTAAGGCATCATTGGA | 83 | ref gene |
| *Tbx21* | GGGCTTCCAACAATGTGACC | AGCTGAGTGATCTCTGCGTTC | 193 | CD4 Tcells ILC |
| *Timp1* | GGGGTGTGCACAGTGTTTC | GACCTGATCCGTCCACAAAC | 81 | Gut barrier |
| *Tlr2* | GCATCCGAATTGCATCACCG | ACAGCGTTTGCTGAAGAGGA | 136 | TLR signalling |
| *Tlr3* | GAATCACAATCGCGCACCAA | CCATAGGGACAAAAGTCCCCC | 178 | TLR signalling |
| *Tlr4* | CTCTCATGGCCTCCACTGGT | TTAGGAACTACCTCTATGCAGGGAT | 137 | TLR signalling |
| *Tlr5* | GATGGATGCTGAGTTCCCCC | AAAGGCTATCCTGCCGTCTG | 139 | TLR signalling |
| *Tnfa* | CAAATGGCCTCCCTCTCATCA | TGGGCTACAGGCTTGTCAC | 88 | Innate; M1 Macrophages ; Th1 |
| *Tnfsf15* | CCATCCTCGCAGGACTTAGC | TGCCTCTGGGAGGTGAGTAA | 135 | Th1 |
| *Tuba* | TGTCCTGGACAGGATTCGC | CTCCATCAGCAGGGAGGTG | 115 | Ref gene |
| *Zbtb16* | GCACTACAGGGTTCACACAGG | CACCGTTGTGTGTTCTCAGG | 107 | NK;ILC |
